# Quantitative intravital imaging reveals *in vivo* dynamics of physiological-stress induced mitophagy

**DOI:** 10.1101/2020.03.26.010405

**Authors:** Paul J. Wrighton, Arkadi Shwartz, Jin-Mi Heo, Eleanor D. Quenzer, Kyle A. LaBella, J. Wade Harper, Wolfram Goessling

## Abstract

Mitophagy, the selective recycling of mitochondria through autophagy, is a crucial metabolic process induced by cellular stress, and defects are linked to aging, sarcopenia, and neurodegenerative diseases. To therapeutically target mitophagy, the fundamental *in vivo* dynamics and molecular mechanisms must be fully understood. Here, we generated mitophagy biosensor zebrafish lines expressing mitochondrially targeted, pH-sensitive, fluorescent probes mito-Keima and mito-EGFP-mCherry and used quantitative intravital imaging to illuminate mitophagy during physiological stresses—embryonic development, fasting and hypoxia. In fasted muscle, volumetric mitolysosome size analyses documented organelle stress-response dynamics, and time-lapse imaging revealed mitochondrial filaments undergo piecemeal fragmentation and recycling rather than the wholesale turnover observed in cultured cells. Hypoxia-inducible factor (Hif) pathway activation through physiological hypoxia or chemical or genetic modulation also provoked mitophagy. Intriguingly, mutation of a single mitophagy receptor *bnip3* prevented this effect, whereas disruption of other putative hypoxia-associated mitophagy genes *bnip3la* (nix), *fundc1, pink1* or *prkn* (Parkin) had no effect. This *in vivo* imaging study establishes fundamental dynamics of fasting-induced mitophagy and identifies *bnip3* as the master regulator of Hif-induced mitophagy in vertebrate muscle.

---

Mitophagy is the removal of damaged mitochondria via autophagy, the lysosomal pathway responsible for degrading and recycling cellular components. It prevents cellular damage and apoptosis (Youle, 2019), and defective mitophagy has been linked to both acute and chronic disease states, including muscular dysfunction (Sandri, 2013), neurodegenerative diseases (Pickrell & Youle, 2015), and drug-induced liver injury (Williams *et al*, 2015; Wrighton *et al*, 2019). Occurrence of baseline mitophagy has also been detected in a variety of cells, especially those with high metabolic rates (McWilliams *et al*, 2016, 2018; Sun *et al*, 2015). However, the full extent of basal mitophagy is incompletely studied, especially during embryonic development. This highlights the need for the development of accurate systems to monitor mitophagy *in situ* that can provide novel mechanistic insight into system dynamics and activation triggers and could enable effective targeting for interventional or therapeutic purposes.

Mitophagy occurs following mitochondrial damage when the autophagosome, a specialized double-membrane organelle, is recruited to the damaged mitochondrial fragment, and then fuses with a lysosome, initiating cargo degradation (Anding & Baehrecke, 2017). However, the organelle dynamics and interactions between mitochondria, autophagosomes, autolysosomes, and lysosomes in response to physiological stresses are underexplored *in vivo*. Fasting is well documented to induce lysosome biogenesis and autophagy (Settembre *et al*, 2013; Raefsky & Mattson, 2017; Preidis *et al*, 2017; Sebastián & Zorzano, 2020). In contrast, dynamic changes in mitolysosome (lysosomes containing mitochondria for degradation) size or number in response to increasing stress *in vivo* are unknown. Further, the temporal dynamics of mitochondrial recycling and trafficking to autolysosomes have not been verified via intravital imaging in any live vertebrate organism.

Multiple molecular pathways govern mitophagy depending on the cell type and nature of stress. PINK1 and PARKIN, genes with mutant alleles linked to Parkinson’s Disease, have been shown to facilitate mitophagy following mitochondrial membrane depolarization, and most molecular mechanisms governing mitophagy were initially described in cell lines overexpressing PARKIN in response to non-physiological stressors, such as exposure to mitochondrial uncouplers (Pickrell & Youle, 2015). In contrast, hypoxia is a physiological stress that induces mitophagy in cultured mouse embryonic fibroblasts (MEFs) (Zhang *et al*, 2008) as well as murine liver (Sun *et al*, 2015), muscle (Zhang *et al*, 2008), and bone marrow (Zhang *et al*, 2016b). The mechanisms governing hypoxia-induced mitophagy remain to be fully interrogated with conflicting evidence pointing towards activation of specific mitophagy receptors such as BNIP3, FUNDC1, or NIX (Palikaras *et al*, 2018; Zhang *et al*, 2016b) and either dependent (Kim *et al*, 2019; Zhang *et al*, 2016a) or independent (Lee *et al*, 2018) of the PINK1-PARKIN pathway. Thus, genetic epistasis analyses using *in vivo* models to dissect which mitophagy receptors or pathways are active in different tissues and under different stresses will further elucidate this process.

Here, we generate mitophagy biosensor zebrafish and define mitochondrial dynamics during vertebrate organ development and complex physiological stresses. Zebrafish develop completely externally and demonstrate a high degree of conservation to mammalian tissue organization, organ development, and molecular signaling (Cox & Goessling, 2015). This vertebrate model system offers the unique ability to illuminate and measure *in vivo* mitophagy trafficking and subcellular dynamics. In this study, we discover widespread basal mitophagy in many organs during development. Further, using time-lapse imaging, we illuminate mitophagy dynamics *in vivo*, directly observing the trafficking of mitochondrial fragments to autolysosomes as it occurs *in situ* in fasted muscle. Using volumetric imaging, we then demonstrate that physiological hypoxia or chemical or genetic activation of the Hypoxia inducible factor (Hif) pathway induce mitophagy in skeletal muscle. Importantly, using precise genetic loss-of-function analyses, we demonstrate that *bnip3* is singularly necessary for Hif-induced mitophagy in muscle independent of other reported hypoxia-associated mitophagy receptors and independent of the *pink1-prkn* (Parkin) pathway. In sum, we established fundamental *in vivo* dynamics of fasting-induced mitophagy and elucidated the genetic and molecular mechanisms governing hypoxia-induced mitophagy.

## RESULTS

### Biosensor zebrafish lines report *bona fide* mitophagy during development

Recently, mitochondria-targeted, pH-sensitive fluorescent protein constructs—mito-mKeima (mito-Keima) (Katayama *et al*, 2011) and mito-EGFP-mCherry (mito-GR; Green Red) (Rojansky *et al*, 2016)—have been developed as sensitive readouts of mitophagy and incorporated into cell culture models and model organisms (Lee *et al*, 2018; McWilliams *et al*, 2016; Sun *et al*, 2015; Ordureau *et al*, 2020). Keima is a fluorescent, red light-emitting protein with a pH-sensitive excitation spectrum, enabling differentiation between Keima in a slightly basic pH environment, such as the healthy mitochondrial matrix (Porcelli *et al*, 2005), or an acidic environment, such as an autolysosome (Katayama *et al*, 2011). Whereas EGFP is sensitive to acid quenching and lysosomal degradation, mCherry and Keima are resistant to lysosomal degradation, allowing for signal accumulation, which is an indicator of the extent of mitophagic flux (Katayama *et al*, 2008).

Zebrafish lines *Tg(ubi:mito-Keima)* and *Tg(ubi:mito-GR)* were generated to express mito-Keima or mito-GR under the control of the ubiquitous *ubiquitin* promoter (Kwan *et al*, 2007; Mosimann *et al*, 2011). Importantly, these reporters are targeted to the mitochondrial matrix because reporters targeted to other mitochondrial locations can lead to false positive signals (Katayama *et al*, 2020). Healthy mitochondria maintain a pH 7.8 matrix (Porcelli *et al*, 2005) and can be distinguished by either a high Keima440/Keima561 ratio (Keima signal when excited by a 440 nm laser divided by Keima signal when excited by a 561 nm laser) or presence of both EGFP and mCherry signal. In contrast, mitochondria undergoing degradation exist in acidic autolysosomes and are distinguished by a high Keima561/Keima440 ratio for mito-Keima or presence of mCherry signal alone for mito-GR.

Ubi:mito-Keima and ubi:mito-GR transgenic zebrafish demonstrated no physiological abnormalities to suggest fluorophore toxicity and were generally analyzed as larvae from 1 to 8 days post fertilization (dpf). To test whether the ubi:mito-Keima construct faithfully reports mitophagy *in vivo*, we exposed ubi:mito-Keima larvae to MitoTracker Green dye, which labels all mitochondria regardless of polarization state, and found that both Keima440^high^ puncta and Keima561^high^ puncta colocalized with MitoTracker Green signal in skin cells (Supplemental Figure S1A). Thus, mito-Keima is correctly localized to mitochondria even after transition to a low-pH environment. In skeletal muscle cells, LysoTracker dye, which accumulates in lysosomes, colocalized with Keima561^high^ puncta, but not with Keima440^high^ puncta (Supplemental Figure S1B). Thus, areas with a high Keima561/Keima440 signal ratio represent mitolysosomes. Further, we crossed ubi:mito-Keima fish with *Tg(CMV:EGFP-Lc3)* fish, which label autophagosomes and autolysosomes with EGFP (He *et al*, 2009). In skeletal muscle, Keima561^high^ fluorescence colocalized with the EGFP-Lc3 signal, indicating that to induce degradation, mitochondrial fragments are shuttled to low-pH, Lc3^+^ autolysosomes (Supplemental Figure S1C). Similarly, we incubated ubi:mito-GR embryos with LysoTracker deep red and observed colocalization with mCherry^+^;GFP^-^ puncta (Supplemental Figure S1D). To ensure macroautophagy was necessary for the reporter signal, larvae were subjected to morpholino (MO)-mediated knockdown of the essential autophagy gene *atg5*. This induced cardiac and central nervous system defects in both ubi:mitoKeima and ubi:mito-GR embryos as previously described (Lee *et al*, 2016, 2014). However, notochords developed similarly at 48 hpf and displayed greatly reduced Keima561^high^ (Figure 1A) and mCherry^+^;EGFP^-^ signals (Figure 1B). Thus, both biosensor lines faithfully differentiate between healthy mitochondria and those undergoing mitophagy.

**Figure 1.**
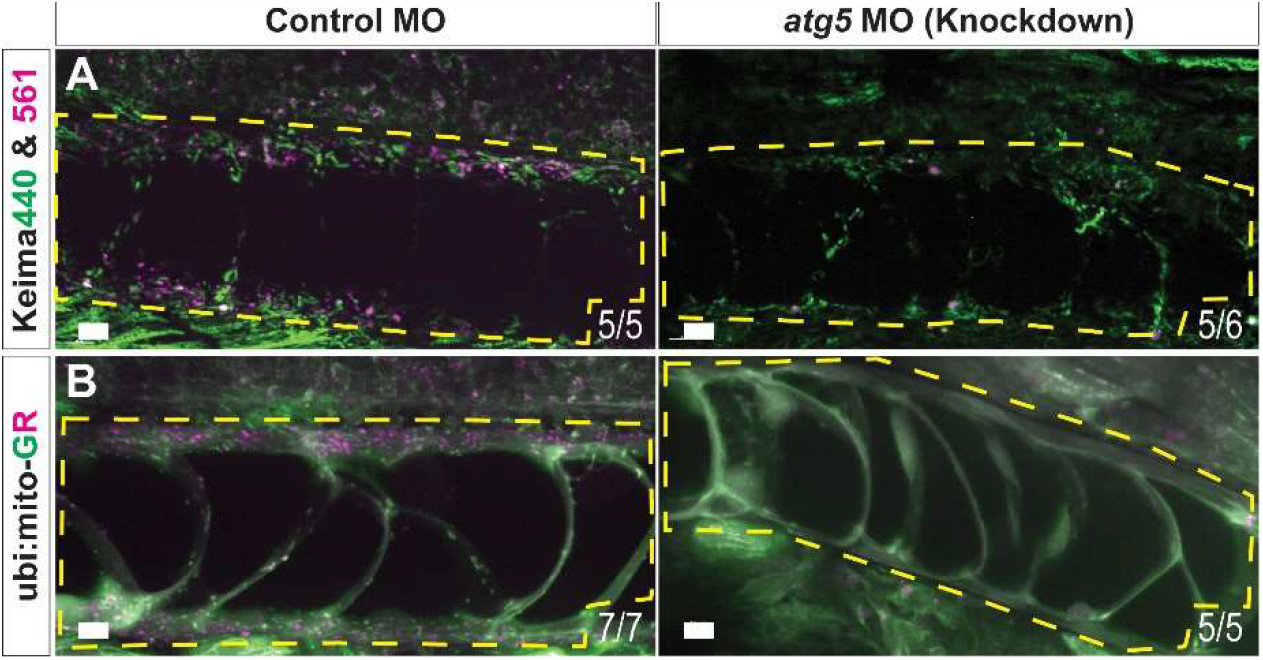
mito-Kiema and mito-GR zebrafish report *bona fide* mitophagy. (A) Maximum intensity projection (MIP) confocal images of 48 hpf ubi:mitoKeima and ubi:mito-GR larvae injected with control or atg5-targeting morpholinos. Yellow dashed boxes highlight the notochord. Control MO-injected larvae have prominent levels of mitophagy (indicated by magenta puncta), whereas *atg5* MO-injected larvae have reduced mitophagy in the notochord. The numbers in the bottom right corners indicate the proportion of larvae observed with similar phenotypes. Scale bars, 10 μm.

While basal mitophagy has been observed in many highly metabolic adult tissues (McWilliams *et al*, 2016), much less is known about mitophagy during vertebrate development. We utilized the mito-GR line to delineate basal levels of mitophagy during organogenesis and observed basal mitophagy in several larval tissues, indicated by dense mCherry^+^;EGFP^-^ signal. Neural tissues including the hindbrain, olfactory placode, trigeminal ganglion, retina, spinal cord, and neuromasts had high levels of basal mitophagy. The developing notochord, vasculature, heart (ventricle), kidney, and liver all displayed high levels of basal mitophagy, and the pancreas displayed moderate levels (Figure 2A). Prior to 4 dpf, larval skeletal muscle exhibited minimal mitophagy. However, after 5 dpf, larvae exhibited a dramatic increase in mitophagy (Figure 2B, arrows). The timing of the onset of skeletal muscle mitophagy coincided with the stage during which zebrafish larvae must transition from subsisting on yolk sac deposited nutrients to external food sources absorbed through their gut (Wilson, 2012) and suggests this observation may be caused by nutrient-deprivation. These results reveal widespread basal mitophagy throughout development and identify zebrafish skeletal muscle as uniquely suited to investigate stress-induced mitophagy *in vivo*.

**Figure 2.**
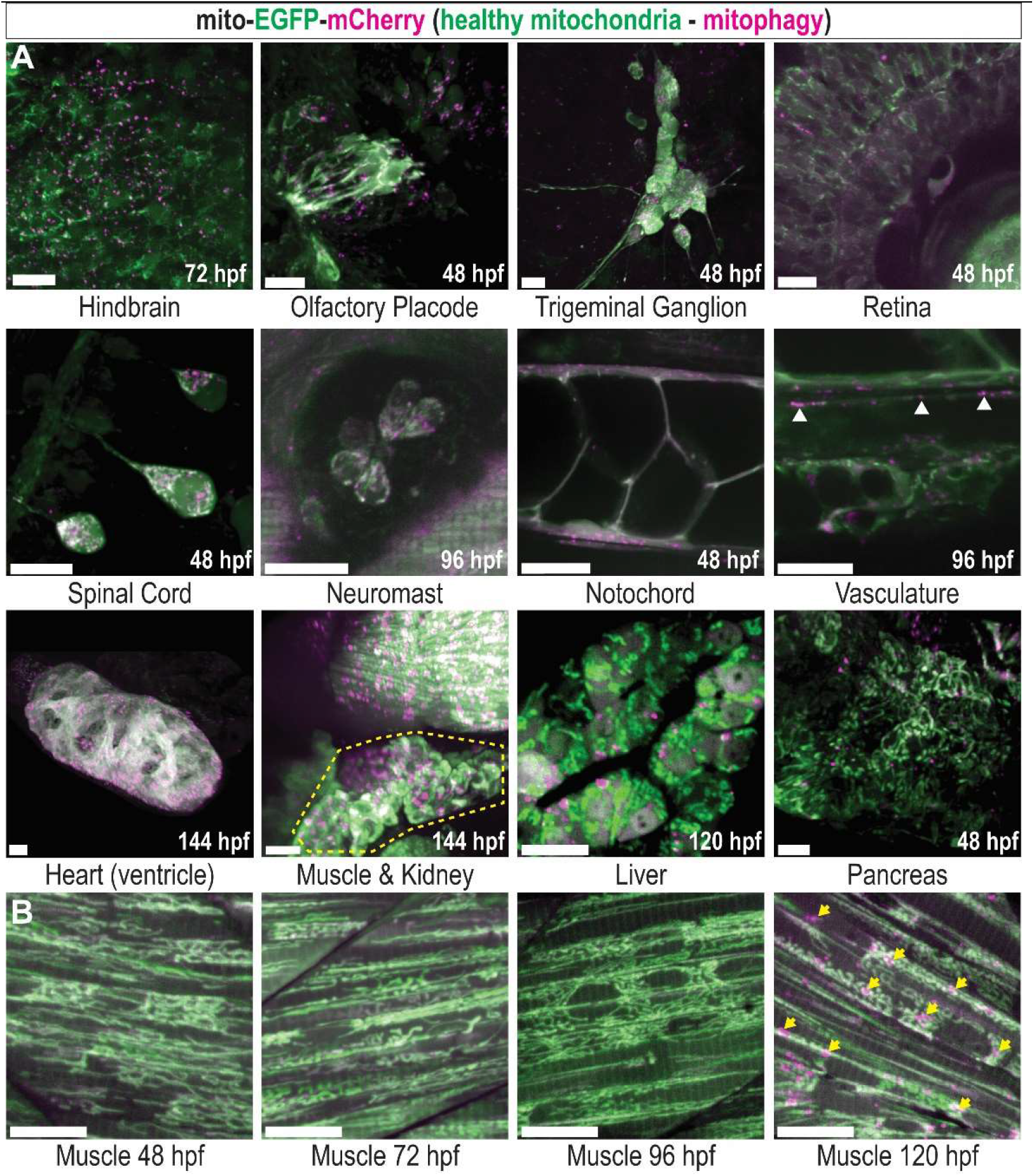
Basal mitophagy is widespread in developing zebrafish tissues. (A) MIP confocal images of mito-GR zebrafish larvae at the indicated developmental time points, displaying various developing organs and tissues. Green indicates EGFP^high^ mitochondria, which are healthy, and magenta indicates mCherry^high^ puncta, representing mitolysosomes (mitophagy). White arrowheads show mitophagy occurring in the vasculature. Yellow outline highlights the developing kidney. (B) Developmental time series of mito-GR skeletal muscle in the tail. Mitophagy is low at 48, 72, and 96 hpf but high at 120 hpf (arrows). All images are of live ubi:mito-GR larvae except the liver image, which is from a hepatocyte-specific fabp10a:mito-GR larva, and the heart image, which is a fixed wholemount ubi:mito-GR larva. Scale bars, 15 μm.

### Fasting induces quantifiable mitophagy and reveals mitolysosome dynamics

Skeletal muscle is a mitochondria-rich, highly metabolic tissue, which responds to nutrient deprivation by inducing autophagy (Mammucari *et al*, 2007), yet the regulation and dynamics of muscle mitophagy *in vivo* are not fully investigated. To precisely quantify mitophagy levels, we developed volumetric and ratiometric image acquisition and analysis methods to rapidly quantify the number and size of mitolysosomes within 3D tissue volumes. Spinning disc confocal fluorescence imaging enabled rapid generation of 3D images of zebrafish muscle spanning >6 myotomes more than 150 μm deep (Figure 3A) at mitochondrial resolution (Figure 3A’; inset of Figure 3A). The Keima561/Keima440 signal ratio allows isolation of mitolysosomes (Figure 3A’’). the number and volume of which can be measured by applying an isosurface rendering algorithm. Normalizing either mitolysosome number or volume to the total tissue volume yields two mitophagy quantification metrics (Figure 3B and Movie S1). This process is also applicable to mito-GR, however at greater imaging depths the ubi:mito-Keima signal decayed more consistently and provided higher signal-to-noise. Thus, we applied this method to quantify fasting-induced mitophagy in ubi:mito-Keima zebrafish in two modes: mitolysosome density, defined as the number of mitolysosomes per tissue volume, and mitophagy index, defined as the proportion of total tissue volume occupied by mitolysosomes.

**Figure 3.**
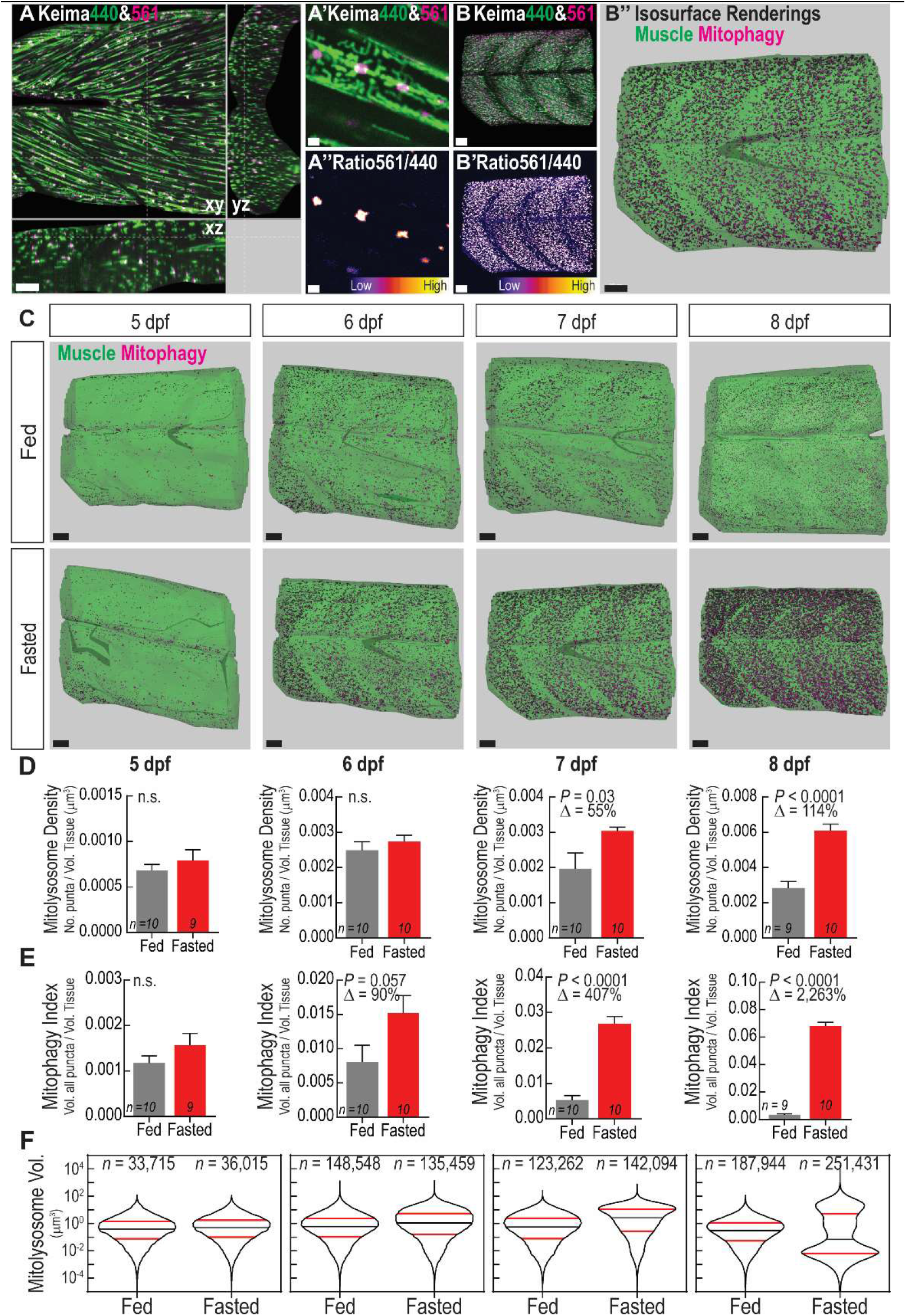
Fasting induces mitophagy in skeletal muscle and volumetric imaging reveals mitolysosome size dynamics. (A) Confocal microscopy cross-sectional image of 6 dpf ubi:mito-Keima zebrafish skeletal muscle demonstrates the typical imaging volume achieved for analyses. Scale, 100 μm. (A’) Zoomed inset of (A) image demonstrates mitochondrial resolution achieved for all images and (A’’) ratiometric images used to segment mitolysosomes. Scale, 3 μm. (B,B’,B”) Representation of the image analysis process. (B) 3D reconstruction of the large volume image showing Keima440 and Keima561 signal throughout greater than 6 myotomes. Scale, 30 μm. (B’) Ratiometric image isolates Keima561^high^ mitolysosomes. Scale, 30 μm. (B”) Isosurface renderings of muscle tissue volume (green) and the mitolysosomes (magenta) allow for volumetric analysis of mitophagy. Scale, 30 μm. (C) Isosurface renderings of muscle volume and mitolysosomes in 5-8 dpf zebrafish either fed or fasted. Green is the surface rendering of the muscle tissue, and magenta are surface renderings for individual mitolysosomes. Scale, 30 μm. (D) Quantification of Mitolysosome Density defined as the number of mitolysosomes divided by the total tissue volume. (E) Quantification of Mitophagy Index defined as the sum of the volumes of all mitolysosomes divided by the total tissue volume. (D,E) Data shown are mean + SEM. *P*, unpaired, two-tailed Student’s t test. n.s., not significant (*P*>0.05). Δ, percentage change of experimental cohort to control cohort. *n* indicates number of larvae in each cohort. (F) Violin plots demonstrating volume (μm^3^) distribution of individual mitolysosomes from all fed or fasted zebrafish in each cohort. Significant changes in the size distributions are evident at every time point (see Figure S2). 50^th^ percentile - black line; 25^th^ and 75^th^ percentiles - red lines. *n*, number of individual mitolysosomes combined from all larvae.

At 5 dpf quantifiable mitophagy was observed in both fed and fasted zebrafish muscle, but there was not a significant difference in the extent measured by mitolysosome density (Figure 3D) or mitophagy index (Figure 3E). However, mitophagy index was a more sensitive readout for mitophagy. Mitolysosome density was not significantly different between fed and fasted larvae until after 7 dpf, whereas the volumetric mitophagy index revealed increases by 6 dpf. Thus, mitophagy in skeletal muscle occurs after yolk depletion, but is abrogated by feeding.

To understand mitolysosome size dynamics during fasting, the volume distributions of individual mitolysosomes were assessed. In fasted skeletal muscle, the early response from 5-7 dpf was an increase in the volume distribution of mitolysosomes. Both fed and fasted fish had a monomodal mitolysosome volume distribution with the fasted fish displaying an increase in 25^th^, 50^th^, and 75^th^ percentile mitolysosome volume (Figure 3F and Supplemental Figure S2). However, by 8 dpf, when fasting entered a more advanced stage, mitolysosome size distribution became bimodal, with an increase in both large and small mitolysosomes (Figure 3F). Surprisingly, this result is unique to mitolysosome dynamics. Although the size distribution of all LysoTracker^+^ lysosomes showed increased size and number in fasted larvae over time, very small lysosomes were present at all stages (Supplemental Figure S3). Together, these results reveal that the early fasting-induced mitophagy response primarily utilizes and expands existing autolysosomes within the cell as evident by an increase in mitophagy index and shift in the size distribution prior to significant changes in the number of mitolysosomes (mitolysosome density). During advanced fasting stages, however, the existing mitolysosomes become large, and new, small mitolysosomes appear. Taken together, these data validate the volumetric mitophagy quantification method and reveal novel mitolysosome size changes as a response to prolonged nutrient deprivation.

To determine the origin of the newly appearing small mitolysosomes found in advanced fasting stages, we performed rapid, continuous time-lapse analysis on a 7 dpf fasted mito-Keima larva (Movie S2). Application of an 3D-object tracking algorithm enabled identification of *de novo* mitophagy and mitolysosome fission events (Figure 4A). Newly appearing Keima561^high^ puncta were identified, and each event was binned into one of several categories. If the Keima561^high^ punctum stemmed from a larger Keima440^high^ mitochondrion, it was considered “new mitophagy”. If the punctum broke away from a different, larger Keima561^high^ mitolysosome, it was considered a “split mitolysosome”. If the punctum migrated into the imaging volume near the edges, it was considered “from outside” (Figure 4B). Out of 381 Keima561^high^ objects tracked for 28 s, 325 were stable from the beginning to the end of the imaging period, 11 migrated from outside, only 3 were new mitophagy (Figure 4C, * and yellow surface object), and 37 puncta split from larger mitolysosomes (Figure 4D, *). Thus, *de novo* mitophagy rapidly occurs during this stage of fasting and accounts for some of the new, small mitolysosomes, but mitolysosome fission is the predominant source of the small mitolysosomes, which appear during advanced fasting.

**Figure 4.**
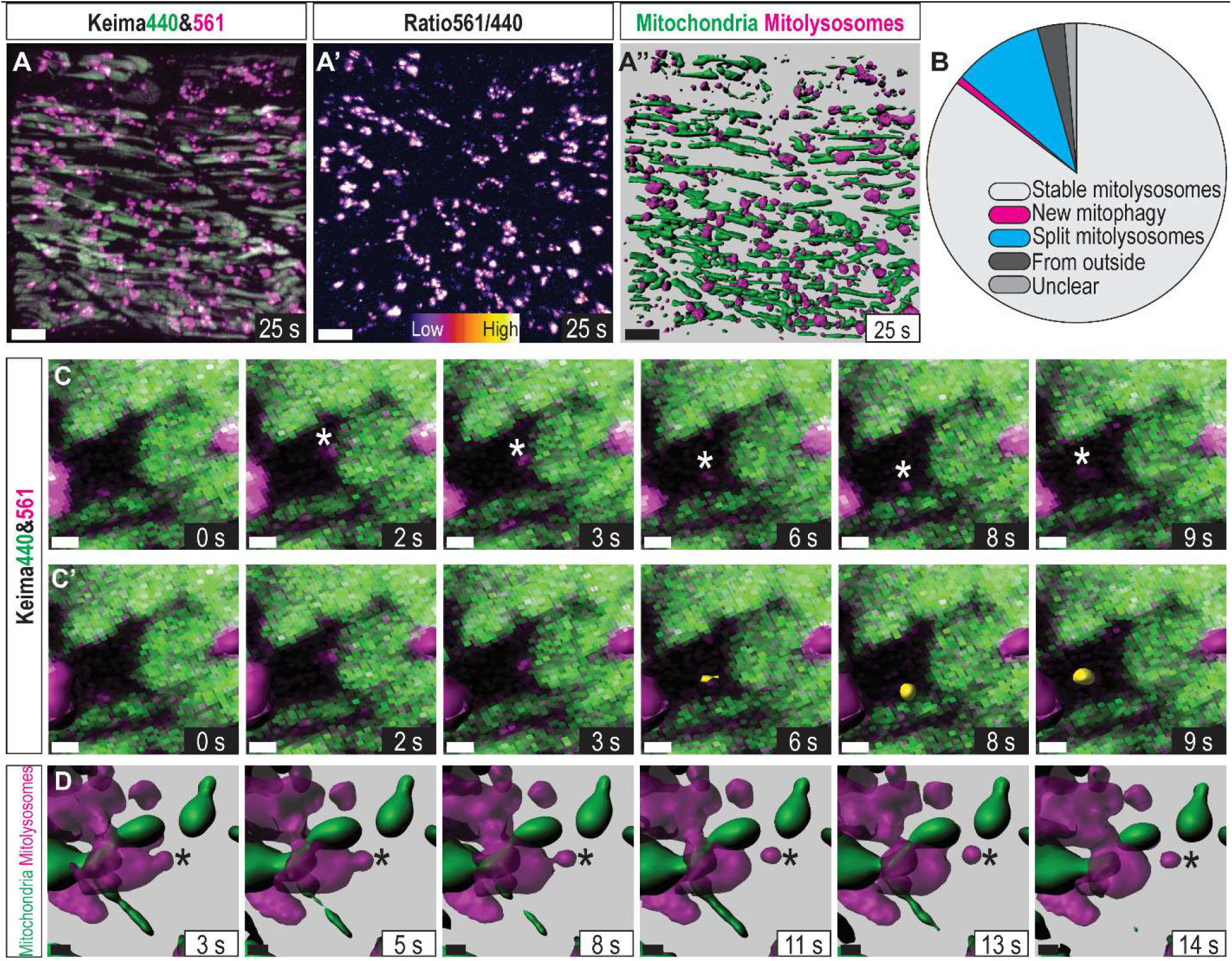
*In vivo* mitophagy dynamics: mitophagy kinetics and mitolysosome fission. (A) Confocal 3D projection of the imaging volume in Movie S2. Images of skeletal muscle in a 7 dpf larva during advanced nutrient deprivation. Images were continuously acquired, finishing each volume approximately once per second and tracking skeletal muscle mitochondria in ~2 cell-deep volume for 28 s. (A’) Ratiometric image isolating mitolysosomes. (A’’) Isosurface rendering of mitochondria (green) and mitolysosomes (magenta) used to track stable mitochondria, ongoing mitophagy, and mitolysosome fission. Scale, 10 μm. (B) Classification of each Keima561^high^ object revealing that approximately 85% were stable, 1% were newly formed, and 10% split from existing, larger mitolysosomes within the 28 s timeframe. The remaining mitolysosomes were pre-existing or translocated from outside the imaging volume. (C) Zoomed timeseries of a representative “new mitophagy” event marked with (*). (C’) Same as (C) with isosurface objects shown for stable mitolysosomes (magenta) and the new Keima561^high^ object (yellow). Scale, 1 μm. (D) Isosurface representation of a splitting mitolysosome (*). Scale, 1 μm.

### Fasting-induced mitophagy occurs piece-by-piece

Because the mitophagy process has not been analyzed by time-lapse imaging in any live vertebrate, we next investigated how GFP-Lc3-labeled organelles interact with mitochondria *in vivo*. By performing time-lapse analysis in *Tg(CMV:GFP-Lc3)^zf155^;(ubi:mito-Keima)* larvae we could directly observe interactions between Lc3-labeled organelles (autophagosomes and autolysosomes) and mitochondrial fragments. Multiple mechanisms have been observed in cells cultured *in vitro* by which a mitochondrion is sequestered by the autophagy machinery. In the “wholesale” mode, after a mitochondrial filament reaches a damage threshold, mitochondrial membrane depolarization and extensive fragmentation occurs, which is followed by recruitment of the autophagy machinery to degrade all of the fragments. In the “piecemeal” mode, the damaged part of a mitochondrial filament is sequestered, and the autophagy machinery is localized to that point, which undergoes fission and degradation, leaving the filament largely intact (Youle, 2019; Yamashita *et al*, 2016; Burman *et al*, 2017). It is unknown whether *in vivo* mitophagy occurs via a wholesale or piecemeal mode. We performed time-lapse analysis in fasted muscle with ~1 min/frame time resolution, and isosurface renderings identified and tracked the different organelle structures in three dimensions. Interestingly, we never observed massive mitochondrial fragmentation followed by mitophagy, indicating the wholesale mode of mitophagy is not active in this context. Mitophagy events (transition from Keima440^high^ to Keima561^high^) occurred at discrete contact points between Lc3 and filamentous mitochondria (Figures 5A and 5A’, * and Movie 3), indicative of piecemeal mitophagy.

**Figure 5.**
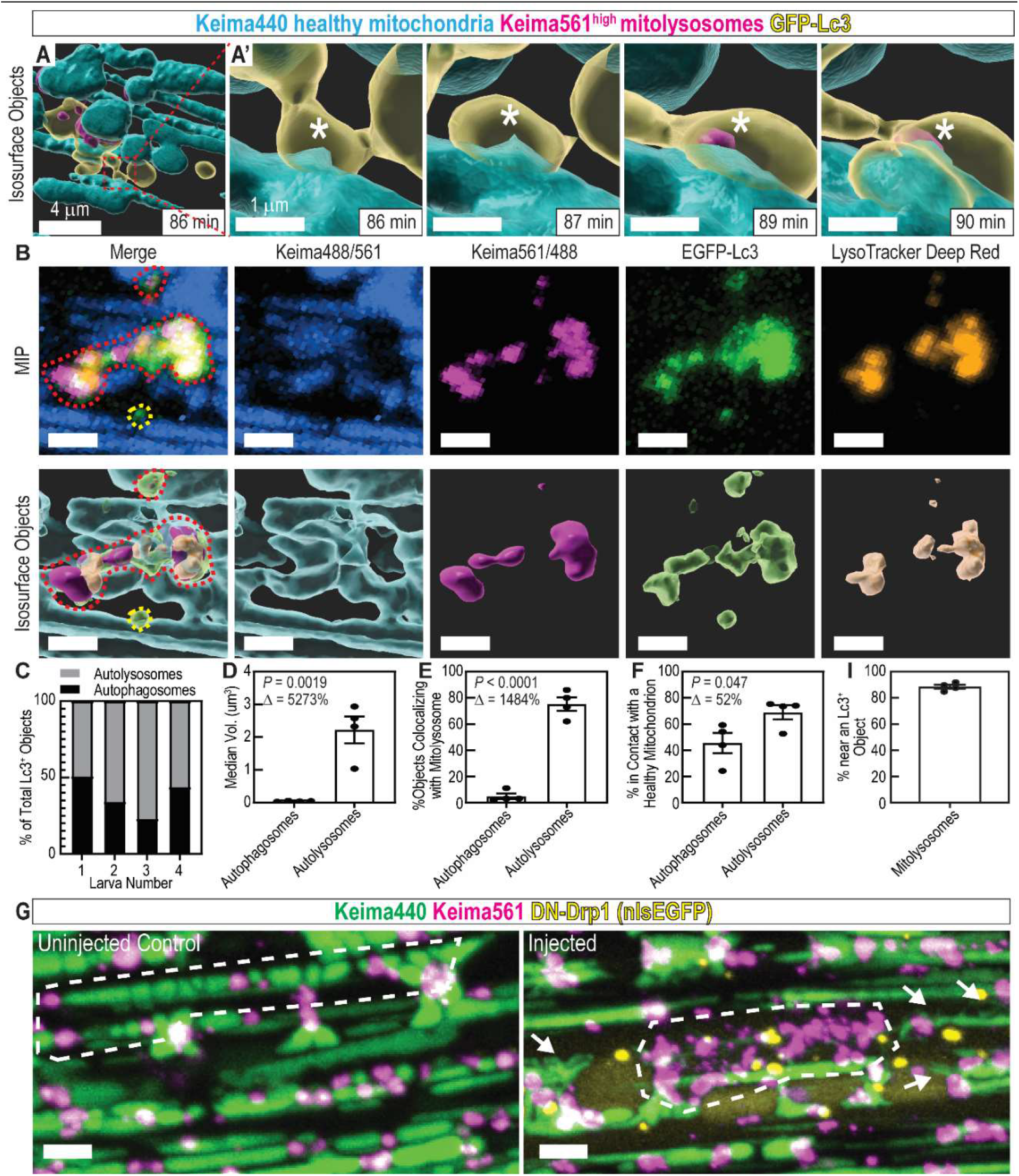
Mitophagy trafficking dynamics: mitophagy occurs piecemeal. (A) Isosurface object projections of confocal time-lapse imaging of fasted skeletal muscle in 5 dpf ubi:mito-Keima larva. Scale, 4 μm. (A’) Inset of (A); mitophagy occurs piecemeal. An EGFP-Lc3^+^ surface is interacting with a Keima440^high^ filamentous mitochondrion (86 min; *). Then, a small portion of the longer filamentous mitochondrion becomes Keima561^high^, indicating pH change and mitophagy (89 min; *). Scale, 1 μm. (B) MIP and isosurface object representations displaying healthy mitochondria (Keima488/561), mitolysosomes (Keima561/488), EGFP-Lc3^+^ objects, and LysoTracker Deep Red^+^ objects. Red outline highlights an autolysosome containing degraded mitochondria. Yellow outline highlights an autophagosome containing no mitochondria. Scale, 2 μm. (C) Classification of EGFP-Lc3^+^ objects as autophagosomes or autolysosomes. Those in contact with LysoTracker^+^ objects are autolysosomes, and the remaining are autophagosomes. (D) Volume quantification of autophagosomes and autolysosomes. (E) Colocalization analysis between mitolysosomes (Keima561^high^ objects) and autophagosomes or autolysosomes. (F) Colocalization analysis between healthy mitochondria and autophagosomes or autolysosomes. (D-F,I) Data shown are mean ± SEM. P, unpaired, two-tailed Student’s t test. Δ, percentage change between autophagosomes and autolysosomes. n = 4 individual 6 dpf larvae. (G) MIP of 7 dpf ubi:mito-Keima larvae injected with an inducible DN-Drp1 construct. Compare the piecemeal mitophagy pattern in the control to the wholesale mitophagy pattern in the injected (white dotted outlines). Mitochondrial morphology is also altered in DN-Drp1-injected larvae (arrows). Scale, 4 μm. (I) Colocalization analysis between mitolysosomes and Lc3^+^ objects.

Intriguingly, the Keima561^high^ mitolysosomes colocalized extensively with EGFP-Lc3 signal, indicating that EGFP-Lc3 labels both autophagosomes and autolysosomes in this system (Supplemental Figure S1C). To differentiate acidified autolysosomes from autophagosomes, *Tg(CMV:GFP-Lc3)^zf155^;(ubi:mito-Keima)* larvae were stained with LysoTracker Deep Red and object-based colocalization image analysis was performed. This analysis required activation of Keima with a 488 nm laser, causing additional overlap between the neutral/basic and acidic Keima signals. To accommodate for this, we defined healthy mitochondria by the Keima488/561 signal ratio and generated surface objects for healthy mitochondria, mitolysosomes, EGFP-Lc3^+^ objects, and LysoTracker^+^ objects (Figure 5B). First, object-based colocalization was employed to determine which EGFP-Lc3^+^ objects were autolysosomes—in contact with a LysoTracker^+^ object—or autophagosomes—not in contact with a LysoTracker^+^ object (Figure 5C). Autolysosomes were substantially larger in volume than autophagosomes (Figure 5D). Remarkably, nearly all autolysosomes contained Keima561/488^high^ objects, classifying them as automitolysosomes, whereas nearly all autophagosomes did not (Figure 5E). Approximately 45% of autophagosomes and ~68% of autolysosomes were in direct contact with healthy mitochondria, supporting our observation that piecemeal mitophagy occurred at discrete contact points between EGFP-Lc3^+^ organelles and larger, intact mitochondrial filaments.

Mitochondrial protein aggregates can be cleared via a piecemeal mitophagy mechanism in cultured cells, and knockout of Drp1 impaired mitochondrial fission, leading to wholesale mitophagy (Burman *et al*, 2017). We hypothesized that inhibition of Drp1 could similarly induce a wholesale mitophagy response *in vivo*. Thus, we introduced a heat shock inducible, dominant negative Drp1(K38A)-2A-nlsEGFP (DN-Drp1) construct (Smirnova *et al*, 1998) to ubi:mito-Keima larvae and induced expression during fasting in the resulting mosaic animals. At 7 dpf, cells expressing DN-Drp1 (identified by nuclear EGFP expression) had altered mitochondrial morphology with increased connections between filaments, indicating that expression of DN-Drp1 functionally reduced mitochondrial fission (Figure 5G, arrows). Interestingly, mitophagy was not obviously diminished in these cells compared to neighboring wild type (WT) EGFP^-^ cells, and in some EGFP^+^ cells, Keima561^high^ signal occurred in a pattern indicative of wholesale mitophagy of large mitochondrial filaments (Figure 5G). Together, these results indicate fasting-induced mitophagy in skeletal muscle occurs piecemeal, and that Drp1 inhibition does not prevent mitophagy but can induce wholesale recycling of mitochondrial filaments.

Mitochondrial substructures can also be removed piecemeal by an Lc3-independent mechanism utilizing mitochondria-derived vesicles (MDVs) (Sugiura *et al*, 2014; Cadete *et al*, 2016). 88.5% of mitolysosomes, however, colocalized within 0.5 μm of an Lc3^+^ object (Figure 5I). Thus, in fasted muscle, the MDV pathway may not play a major role in mitochondrial recycling. In sum, mitophagy in fasted skeletal muscle occurs primarily by a piecemeal mechanism utilizing Lc3^+^ organelles.

### Hif activation induces mitophagy

Tissue hypoxia (ischemia) occurs in muscle during in exercise and other physiological and pathophysiological conditions (Leermakers & Gosker, 2016; Lee *et al*, 2019; Ferdinand & Roffe, 2016). This stress leads to production of excess reactive oxygen species, mitochondrial damage, and mitophagy (Kaelin & McKnight, 2013). We used the zebrafish mitophagy reporter system to dissect the effects of hypoxia and activation of the Hif pathway on muscle mitophagy (Figure 6A). To avoid fasting-induced mitophagy as a confounding variable, we examined larvae younger than 5 dpf. To induce physiological hypoxia, zebrafish larvae were raised in hypoxic fish water within a hypoxia chamber (4.7% O_2_) from 77-96 hpf, causing death in 11 of 35 larvae, indicative of the substantial hypoxic stress induced by this approach. Surviving larvae had a significant increase in mitophagy in skeletal muscle (Figure 6B). Next, we tested whether chemical activators of Hif signaling induced mitophagy in skeletal muscle. The PHD inhibitor dimethyloxalylglycine (DMOG) has been validated to induce Hif activation in similarly aged zebrafish larvae (Elks *et al*, 2013; Ivan *et al*, 2001; Iliopoulos *et al*, 1996). Fish exposed to DMOG from 3-4 dpf exhibited robust mitophagy compared to controls (Figure 6C). This result was confirmed by exposing zebrafish to FG-4592 (Roxadustat), a chemically dissimilar PHD inhibitor evaluated for FDA approval, which caused a similar increase in mitophagy (Figure 6D). Genetic knockdown of *vhl* using a previously validated MO (Harris *et al*, 2013) also induced mitophagy at 3 dpf (Figure 6E). To test whether Hif activation alone was sufficient to induce mitophagy in skeletal muscle in a cell autonomous manner, dominant active Hif (*DA-hif1ab*) (Elks *et al*, 2011) and nuclear-localized EGFP were expressed under the control of the muscle-specific *unc45b* promoter (Berger & Currie, 2013). This resulted in mosaic expression of nuclear EGFP, marking the cells expressing DA-hif1ab to be compared to neighboring EGFP^-^, WT cells within the same animal. EGFP^+^ cells had high levels of mitophagy, whereas surrounding EGFP^-^ cells had little-to-no mitophagy similar to uninjected controls (Figure 6F). Together, these results confirm that hypoxia and Hif activation elicit mitophagy in a cell-autonomous fashion.

**Figure 6.**
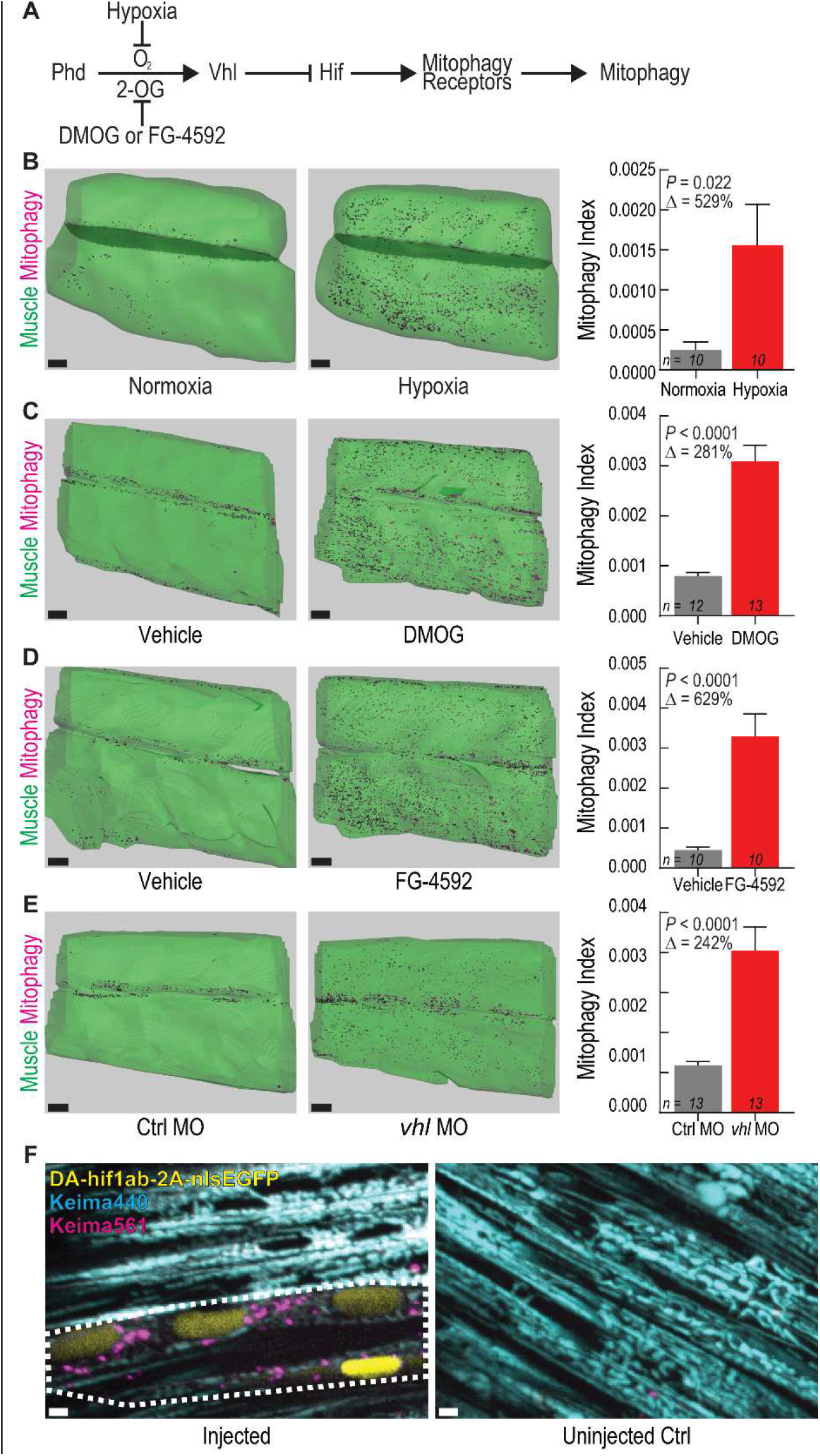
Activating hypoxia-inducible factor (Hif) signaling induces mitophagy in skeletal muscle. (A) Schematic of Hif signaling pathway. 2-OG, 2-oxoglutarate. (B-E) Isosurface renderings and Mitophagy Index quantification in skeletal muscle of 4 dpf ubi:mito-Keima zebrafish exposed to (B) 4.7% oxygen (hypoxia), (C) 60 μM DMOG, or (D) 10 μM FG-4592 from 3-4 dpf. (E), Isosurface renderings and quantification of mitophagy in skeletal muscle of 3 dpf zebrafish injected with control MO (Ctrl) or *vhl* antisense MO. Scale bars, 30 μm. Data shown are mean + SEM. *P*, unpaired, two-tailed Student’s t test. Δ, percentage change of experimental cohort to control cohort. *n* indicates number of larvae in each cohort. (F) Confocal microscopy images of 4 dpf ubi:mito-Keima expressing dominant active (DA) *hif1ab* in a mosaic fashion. Keima440 is cyan. Keima561 is magenta. Nuclear EGFP is yellow and indicates the cells that express *DA-hif1ab*. White dotted line outlines two adjacent myotubes expressing *DA-hif1ab* and undergoing mitophagy. Scale bars, 2 μm.

### Disruption of *bnip3* alone diminishes Hif-induced mitophagy

Next, we used the volumetric *in vivo* mitophagy quantification method and high-efficiency genome editing to dissect the molecular mechanisms required for Hif-induced mitophagy in vertebrate skeletal muscle. As a positive control, we tested whether the mitophagy signal resulting from DMOG exposure was dependent on macroautophagy by assessing dependence on Vps34 function, which is necessary for autophagosome formation and vital for macroautophagy. A cohort of DMOG-exposed larvae (both ubi:mitoKeima and ubi:mito-GR in independent experiments) was treated with the Vps34 inhibitor SAR405 (Ronan *et al*, 2014). Exposure to SAR405 abrogated DMOG-induced mitophagy (Figure 7A and Supplemental Figure S4), indicating that macroautophagy is vital for Hif-induced mitophagy.

**Figure 7.**
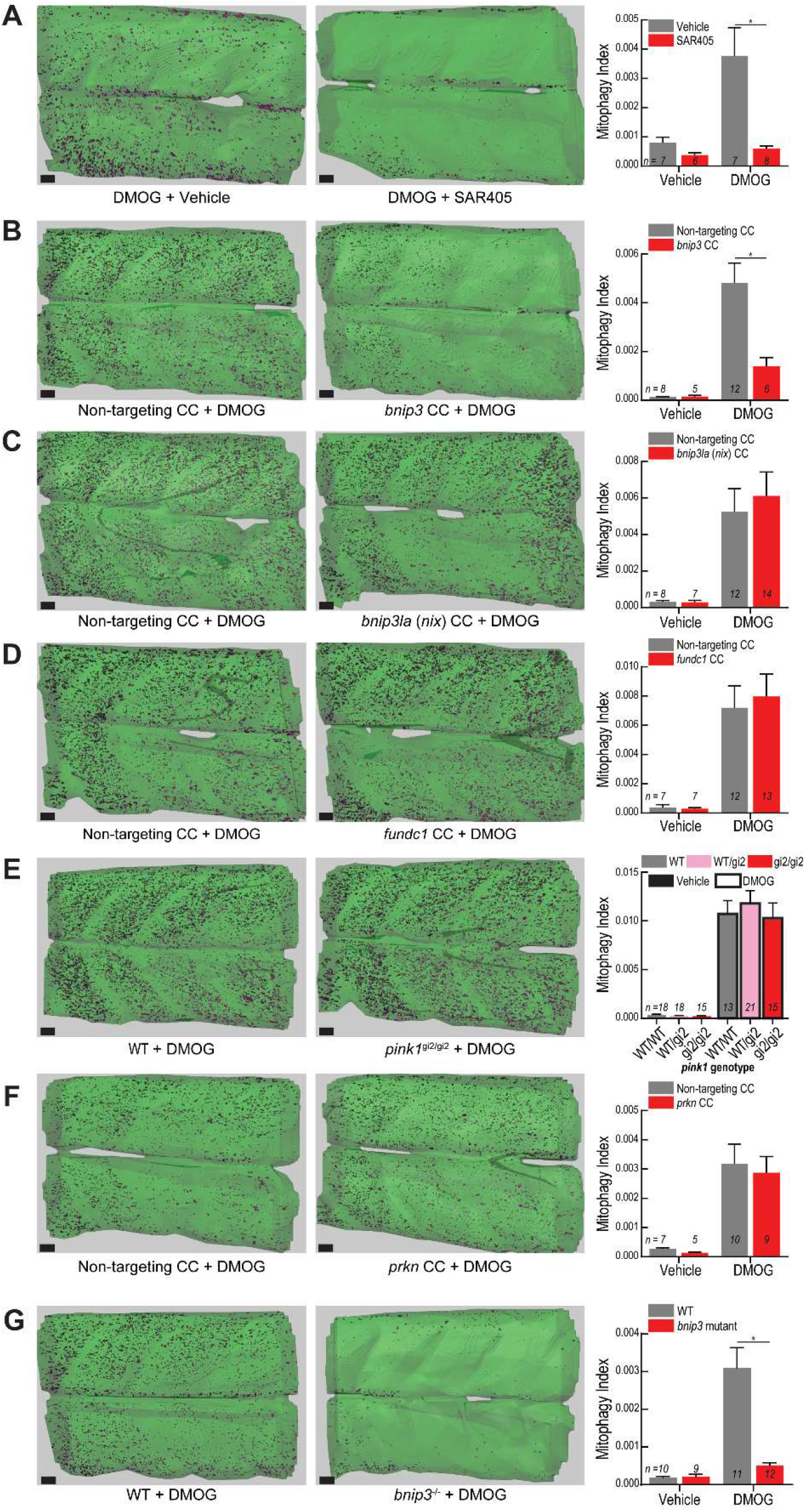
*bnip3* regulates Hif-induced mitophagy in muscle independent of *bnip3la (nix), fundc1, pink1*, and *prkn*. (A) Mitophagy Index quantification of larvae treated with 100 μM DMOG or vehicle (DMSO) and 20 μM SAR405 or vehicle (DMSO). Representative isosurface rendering images show the effect of SAR405 treatment on DMOG-induced mitophagy. (B-D,F) Mitophagy Index quantification and representative isosurface rendering images of DMOG-induced mitophagy in larvae injected with CRISPR cutters (CC) targeting (B) *bnip3*, (C) *bnip3la (nix)*, (D) *fundc1*, or (F) *prkn*. Scale bars, 20 μm. (E) Mitophagy Index quantification and representative isosurface rendering images of DMOG-induced mitophagy in larvae carrying a null *pink1* allele (gi2) and siblings. (G) Mitophagy Index quantification and representative isosurface rendering images of DMOG-induced mitophagy in larvae carrying a 5-bp deletion mutation in the *bnip3* gene. (A-G) Data shown are mean + SEM. \**P*<0.01, two-way ANOVA with Holm-Sidak’s multiple comparisons test. Each comparison between vehicle- and DMOG-treated samples was also significant (*P*<0.05) except in (A) vehicle controls to DMOG+SAR405, (B) vehicle controls to DMOG + *bnip3* CC, and (G) vehicle controls to *bnip3^-/-^* + DMOG. Scale, 20 μm. See Supplemental Figures S4, S6-S8 for replicates and full statistical analyses. *n* indicates number of larvae in each cohort.

HIF-induced mitophagy is thought to rely on the function of specific mitophagy receptors BNIP3, BNIP3L (NIX), and FUNDC1, which contain LC3-interacting regions (LIRs) (Birgisdottir *et al*, 2013; Yoo & Jung, 2018; Hanna *et al*, 2012). Further, the PINK1-PARKIN pathway governs mitochondrial depolarization-induced mitophagy (Pickles *et al*, 2018; Harper *et al*, 2018), but its role in hypoxia-induced mitophagy remains controversial with evidence in other model organisms both for (Kim *et al*, 2019; Zhang *et al*, 2016a) and against (Lee *et al*, 2018) a role. To test the requirement of these factors for Hif-induced mitophagy, we performed a targeted genetic epistasis screen. *bnip3, bnip3la (nix)*, and *fundc1* were disrupted in F0 larvae by CRISPR methodology, an established and rapid method to assess loss of gene function (Supplemental Figure S5) (Burger *et al*, 2016; DiNapoli *et al*, 2020). Surprisingly, larvae injected with bnip3-targeted CRISPR cutter (CC) reagents reliably exhibited dramatic reduction in DMOG-induced mitophagy, indicating that singular disruption of the *bnip3* locus diminishes Hif-induced mitophagy in vertebrate skeletal muscle (Figure 7B and Supplemental Figure S6). Disruption of other receptors implicated in hypoxia-induced mitophagy, specifically *bnip3la (nix)* (Figure 7C) and *fundc1* (Figure 7D), had no effect on DMOG-induced mitophagy. These results indicate that *bnip3*, a single mitophagy receptor, is necessary for the Hif-induced mitophagy response in vertebrate skeletal muscle.

BNIP3 has been shown to interact directly with PINK1, promoting mitophagy via the PINK1-PARKIN pathway (Zhang *et al*, 2016a), and a study in flies showed *Pink1* to be necessary for hypoxia-induced mitophagy (Kim *et al*, 2019). However, there were no significant differences in Hif-induced mitophagy in skeletal muscle between fish homozygous for an established *pink1* null allele (gi2) (Zhang *et al*, 2017) and their WT or heterozygous siblings (Figure 7E and Supplemental Figure S7). Larvae injected with a validated *pink1* MO (Priyadarshini *et al*, 2013) also showed no difference in DMOG-induced mitophagy compared to controls (Supplemental Figure S7). Further, disruption of *prkn* had no effect (Figure 8F). Lastly, we sought to confirm the F0 CRISPR results by generating a stable *bnip3* mutant line, containing a 5-bp frame-shift deletion in exon 2 and an early stop codon (Supplemental Figure S8). Larvae homozygous for the mutant *bnip3* allele had drastically reduced mitophagy capacity compared to WT when challenged with DMOG (Figure 7G and Supplemental Figure S8). In sum, *bnip3*, a single Hif-responsive mitophagy receptor, mediates Hif-induced mitophagy in skeletal muscle, independent of the *pink1-prkn* pathway.

## DISCUSSION

In this study, we reveal fundamental characteristics of *in vivo* mitophagy during normal organ development and complex physiological stresses using mitophagy biosensor zebrafish combined with time-lapse and quantitative imaging. Basal mitophagy occurred in many developing organs. In fasted skeletal muscle, mitolysosomes initially increased in size, followed by a period of concurrent mitophagy and mitolysosome fission. Time-lapse imaging revealed fasting-induced mitophagy to occur *in vivo* via a piecemeal mechanism. Hypoxia and Hif-activation induced mitophagy in skeletal muscle. Hif-induced mitophagy required *bnip3* function, but *pink1, prkn, bnip3la (nix)*, and *fundc1* are dispensable. In sum, these results reveal *in vivo* subcellular dynamics and molecular mechanisms regulating physiological-stress induced mitophagy.

### Many organs exhibit mitophagy during development

Transgenic animals expressing mitophagy reporter constructs have begun to reveal basal mitophagy levels *in vivo*. Murine studies demonstrated that adult tissues with high metabolic demands—brain, liver, kidney, spleen, skeletal muscle—have high basal mitophagy levels (McWilliams *et al*, 2016; Sun *et al*, 2015). Similarly, studies in flies show basal mitophagy in the central nervous system, epidermis, and muscle (Lee *et al*, 2018). The developing mouse heart and kidney display high mitophagy (McWilliams *et al*, 2016), but more extensive developmental analyses have not been performed. Recently, *nipsnap1* mutant zebrafish were found to have decreased basal mitophagy in skin and defects in dopamine neuron development (Abudu *et al*, 2019). Our study detects widespread mitophagy in embryonic and larval zebrafish during the development of many organs, including several previously unrecognized, such as the spinal cord, notochord, vasculature, pancreas, and liver.

### Mitophagy occurs via a piecemeal process *in vivo*

The *in vivo* dynamics of mitophagy have not been previously studied via live imaging. The canonical mitophagy sequence involves mitochondrial fission occurring prior to engulfment by preformed autophagosomes (Anding & Baehrecke, 2017). Conversely, in HeLa cells, mitophagy receptors localized to the point of damage on mitochondria can recruit the ULK1 complex, initiating autophagosome biogenesis locally (Vargas *et al*, 2019) and concurrently with mitochondrial division (Yamashita *et al*, 2016). In live zebrafish skeletal muscle, we found that many autophagosomes and most autolysosomes were in direct contact with intact mitochondrial filaments, indicating that much of autophagosome biogenesis may occur locally, and that much of the fasting-induced autophagy occurring is mitophagy. Time-lapse analysis indicated that mitophagy, as defined by transition from Keima440^high^ to Keima561^high^, occurred near these points in a piecemeal fashion. This also reveals that the process of autophagosome acidification in skeletal muscle is rapid during this advanced stage of fasting. The piecemeal nature of mitophagy in skeletal muscle may be an adaptation necessary to allow for muscle mitochondria to exist as large, fused networks to more efficiently generate ATP and other metabolites (Romanello & Sandri, 2013; Friedman & Nunnari, 2014). Indeed, in mice, skeletal muscle-specific deletion of Opa1, a protein necessary for inner mitochondrial membrane fusion, leads to fragmented mitochondria, muscle degradation, mitochondrial dysfunction, and mitophagy defects (Rodríguez-Nuevo *et al*, 2018). It will be important to determine why mitophagy occurs at specific points along the mitochondrial filament *in vivo*. The topology and shape of mitochondria have been implicated as driving forces for starvation-induced mitophagy in osteosarcoma cells (Zhou *et al*, 2019). Differential segregation and phosphorylation of mitochondrial matrix proteins was also shown to determine mitophagy rates of different mitochondrial proteins (Abeliovich *et al*, 2013; Kolitsida *et al*, 2019). It is exciting to speculate that different mitochondrial proteins could segregate to specific subdomains and dictate the location of mitochondrial fission and mitophagy.

Different types of piecemeal selective autophagy have been discovered. A LIR-containing protein TEX264 was recently demonstrated to be necessary for piecemeal selective autophagy of the ER in HEK293 cells (Abeliovich *et al*, 2013), proving that receptor-mediated selective autophagy can also occur piecemeal. In HeLa cells, a homeostatic piecemeal mitophagy mechanism relies on p62 and LC3C but not PINK1-PARKIN to regulate oxidative phosphorylation (Le Guerroué *et al*, 2017). A recent study explored mitophagy kinetics using mild oxidative stress to induce more physiological levels of mitophagy in cultured rat neurons, suggesting that mild mitochondrial stresses may better model the *in vivo* process (Evans & Holzbaur, 2020). It is possible the wholesale mechanism only occurs in dying cells or those with defects in mitochondrial fission, or that it is primarily an *in vitro* event caused by unphysiologically high mitochondrial stress levels. It will be important for future work to analyze the mitophagy trafficking dynamics and mechanisms *in vivo* in other cell types and under different physiological stresses to uncover endogenous mitophagy kinetics and to determine if every instance of mitophagy occurs via a piecemeal mechanism.

### *bnip3* is required for hypoxia-induced mitophagy

Many genes have been implicated in both ubiquitin-dependent and ubiquitin-independent mitophagy (Rodger *et al*, 2018; Anding & Baehrecke, 2017; Harper *et al*, 2018). These studies, however, were primarily performed in cell culture with limited confirmation of conserved mechanisms *in vivo*. BNIP3, BNIP3L (Nix), and FUNDC1 have all been linked to hypoxia-induced mitophagy and HIF signaling (Yoo & Jung, 2018; Hanna *et al*, 2012). They contain domains to localize them to the mitochondrial outer membrane, and LIRs, which bind to LC3 and recruit autophagosomal membranes (Birgisdottir *et al*, 2013). BNIP3 mediates hypoxia-induced mitophagy in MEFs and murine lung tissue (Zhang *et al*, 2008). In murine skeletal muscle, FoxO signaling causes fasting-induced autophagy via BNIP3, although neither mitophagy nor hypoxia were analyzed (Mammucari *et al*, 2007). BNIP3 also facilitates mitophagy in murine cortical neurons in a stroke model (Shi *et al*, 2014) and in murine hearts and HL1 cardiac muscle cells after myocardial ischemia/reperfusion injury (Hamacher-Brady *et al*, 2007). BNIP3L (NIX) is linked to hypoxia-induced mitophagy in ischemic brain injury and heart ischemia/reperfusion injury in mice (Yuan *et al*, 2017). FUNDC1 is thought to facilitate hypoxia-mediated mitophagy in HeLa cells (Liu *et al*, 2012; Chen *et al*, 2014), several murine tissues including platelets, liver, heart, and muscle (Zhang *et al*, 2016b), and also in cardiac ischemia/reperfusion injury (Zhou *et al*, 2017). In light of these studies, it might be expected that multiple, redundant mitophagy receptors facilitate the hypoxia response in skeletal muscle. In contrast, we found that mutation of the singular receptor *bnip3* prevented Hif-induced mitophagy in skeletal muscle, whereas disruption of *bnip3la (nix)* or *fundc1* had no effect. Our results add mechanistic insight into the tissue- and stress-dependent contexts for vertebrate mitophagy mechanisms.

There is conflicting evidence for how BNIP3 induces mitophagy. In MEFs, BNIP3 functions by facilitating the release of Beclin-1 from Bcl-2, allowing Beclin-1 to initiate autophagy independent of the PINK1-PARKIN pathway (Zhang *et al*, 2008). However, a separate study in MEFs and HEK293 cells showed BNIP3 interacted with and stabilized PINK1, which recruited PARKIN to the mitochondrial outer membrane and initiated PINK1-PARKIN-dependent mitophagy (Zhang *et al*, 2016a). Separately, *Pink1* was shown to be necessary for hypoxia-induced mitophagy in flies (Kim *et al*, 2019). In zebrafish, however, we tested an established mutant *pink1* allele, *pink1* morphants, and *prkn* disruption via CRISPR and found that none of these interventions influenced or prevented Hif-induced mitophagy in zebrafish muscle. This supports the conclusion that in vertebrates, bnip3-induced mitophagy is independent of the Pink1-Parkin ubiquitination system. In flies, *Pink1* mutation causes muscle degeneration (Clark *et al*, 2006; Yang *et al*, 2006; Park *et al*, 2006), and BNIP3 overexpression can rescue the phenotype (Zhang *et al*, 2016a). In humans (Gasser, 2009), monkeys (Yang *et al*, 2019), mice (McWilliams *et al*, 2018), and zebrafish (Zhang *et al*, 2017), there is no evidence of direct muscle degeneration due to PINK1 mutation, indicating that there are important differences between vertebrate and invertebrate mitophagy mechanisms. Because there are few yeast homologs to vertebrate mitophagy receptors (Mao *et al*, 2011) and due to the observed differences in mitophagy mechanisms between vertebrate and invertebrate models, analysis of mitophagy reporter zebrafish provides a unique avenue toward deeper understanding of mitochondrial quality control dynamics and mechanisms. In summary, our study revealed widespread mitophagy during development, fundamental piecemeal mitophagy dynamics during fasting, and that hypoxia-induced mitophagy in muscle is dependent on *bnip3*.

## MATERIALS AND METHODS

### Animal Studies

Zebrafish (*Danio rerio*) were maintained according to institutional animal care and use committee (IACUC-BIDMC #506-2015) protocols. Adult fish and larva were raised at 28.5 °C. All studies were performed in larval zebrafish before sex can be determined, and therefore sex was not taken into consideration. All embryos and larvae were raised and experimentally treated in E3 salt solution. For experiments involving larval feeding, embryos were provided live paramecia and larval AP-100 fish food (Zeigler) *ad libitum*.

### Molecular Cloning and Transgenesis

Transgenic lines *Tg(ubi:mito-Keima,cryaa:Cerulean), Tg(ubi:mito-GR)*, and *Tg(fabp10a:mito-GR)* were generated using *tol2* transgenesis (Mosimann & Zon, 2011; Mosimann *et al*, 2011; Her *et al*, 2003). Plasmids generated or used are listed in the Supplemental Information Resources Table. Middle entry vector pENTR:mito-Keima was generated via PCR using pHAGE mt-Keima IRES Puro as template and pENTR/D-TOPO kit (Invitrogen) following manufacturer’s instructions. A Gateway reaction using LR Clonase II Plus (Invitrogen) generated the pTol2-*ubi:mito-Keima,cryaa:Cerulean* vector following manufacturer’s instructions. Middle entry vector pME-mito-GR was generated using BP Gateway cloning using pCLBW cox8 EGFP mCherry (Rojansky *et al*, 2016) and pDONR221 (Invitrogen) using Gateway BP Clonase II Enzyme (Invitrogen) mix following manufacturer’s instructions. LR Gateway cloning was used to generate pTol2-*ubi:mito-GR*, and pTol2--*2.8fabp10a:mito-GR*. Middle entry vector pENTR-DA-hif1ab(noStop) was generated using PCR to amplify DA-hif1ab minus the Stop codon from a plasmid containing DA-hif1ab (Elks *et al*, 2011), which was a gift from Phil Elks. Middle entry vector pENTR-DN-Drp1(noStop) was generated similarly using a plasmid containing Drp1(K38A) (Smirnova *et al*, 1998). LR Gateway reactions generated the pTol2--*503unc45b:DA-hif1ab-2A-nlsEGFP,cryaa:mCherry*, pTol2-*hsp70l:DN-Drp1-2A-nlsEGFP,cryaa:mCherry*, and pTol2-*hsp70l:mito-Keima,cryaa:mCherry*. vectors following manufacturer’s instructions. *Tol2* mRNA was synthesized using mMessage mMachine SP6 *In vitro* Transcription kit (Invitrogen) following the manufacturer’s instructions. To generate stable transgenic lines, 1-cell stage TU embryos were injected with the transgenesis vector and *tol2* transposase mRNA. Positive embryos were screened either for cyan fluorescent lenses (ubi:mito-Keima) or for EGFP fluorescence (ubi:mito-GR and fabp10a:mito-GR) and raised to adulthood. F0 adults were outcrossed to TL mates, the F1 generation clutches were screened for bright fluorescence, and potential founders were raised. Potential F1 founders were outcrossed again to TU or TL mates and screened for bright fluorescence and 1:1 ratio of WT to transgenic offspring indicative of single copy insertion (Mosimann & Zon, 2011).

### Fasting and Feeding Protocols

Fasted fish were kept in E3 medium cleaned of any dead larvae and shed chorions. Fed controls were provided live paramecia and AP100 dry larval diet (Zeigler) *ad libitum*. For heat shock induction of mito-Keima and DN-Drp1, injected F0 mosaic larvae were placed into a 50 mL conical tube with 35 mL of E3 then incubated for 45 min at 38 °C before being returned to room temperature E3.

### Hypoxia and Chemical Treatments

For MitoTracker analysis, larvae were exposed to 1 μM MitoTracker Green FM (Cell Signaling Technology) overnight in E3, washed in E3 twice, and prepared for imaging. For LysoTracker analysis, larvae were exposed to 1 μM LysoTracker Green DND-26 (Cell Signaling Technology) or LysoTracker Deep Red (Thermo Fisher Scientific) overnight in E3, washed in E3 twice, and prepared for imaging. For all drug treatment experiments, single clutches were used and split between experimental groups. If additional embryos were necessary, two clutches from sibling breeding pairs were combined and mixed to split randomly. For hypoxia experiments, E3 was reduced to approximately 1.7 mg/L O_2_ by bubbling in a mixture of approximately 96% N_2_ and 4% O_2_ for about 10 minutes. The larvae E3 in a 10 cm dish was replaced with about 40 mL of the hypoxic water, and the dish was moved to a hypoxia chamber, which was then flushed with the same mixture of approximately 96% N_2_ and 4% O_2_ for about 10 minutes until an O_2_ sensor (DO_2_10, Extech Instruments) read 4.7%. The chamber was sealed and moved into the 28.5 °C incubator. For chemical hypoxia experiments, larvae were exposed to 60 μM or 100 μM DMOG (Tocris) or 10 μM FG-4592 (Cayman Chemical) in E3 from about 70 hpf to 98 hpf. The exception is the *pink1* MO experiment for which the treatment window was from 48-72 hpf because MO knockdown is most effective during the first 3 dpf before additional cell divisions dilute the MO (Bill *et al*, 2009). For SAR405 experiments, larvae were split into cohorts receiving vehicle (DMSO; Sigma Aldrich) or 100 μM DMOG and then those cohorts were split into cohorts receiving additional vehicle (DMSO) or 10 μM SAR405 (EMD Millipore). The SAR405 treatment experiment and each high-efficiency CRIPSR experiment was replicated.

### Confocal Microscopy

Microscopy was performed using an inverted Ti2 (Nikon) microscope equipped with a Yokogawa CSUW1 spinning disc confocal unit, a CFI Apo LWD Lambda S 40XC WI (1.15 N.A.) objective lens (Nikon), and a Zyla 4.2+ sCMOS camera (ANDOR). Keima images were captured by exciting with 440 nm laser light and 561 laser light with ET Cy3/TRITC emission filter (Chroma) in triggered acquisition mode. Mito-GR images were captured using 488 nm laser light and 561 nm laser light with ET FITC/TRITC dual emission filter (Chroma) in triggered acquisition mode. For MitoTracker Green, LysoTracker Green, EGFP-Lc3, and nlsEGFP imaging, 488 nm excitation was utilized with a 525/36 FITC emission filter (Chroma). For Lysotraker Deep Red, a similar microscope equipped with a 640 nm laser and ET705/72m emission filter (Chroma) was utilized. For images involving both Keima and EGFP signal, the EGFP images were uniformly frame-shifted (Nikon Elements) based on control images to account for the shift caused by using different dichroic mirrors. For the *in vivo* mitophagy dynamics time-lapse imaging (Figure 4 and Movie 2), an iXon Life 888 EMCCD (ANDOR) was used for its increased light sensitivity and frame rate capacity, allowing greater time resolution. Larvae were anesthetized with 0.16 mg/ mL Tricaine-S (Western Chemical, Inc.) and mounted on glass-bottom dishes (MatTek) in 0.8% low-melt point agarose in E3 containing Tricane. After the agar solidified, additional E3 + Tricane was added to keep the gel hydrated during imaging. For skeletal muscle imaging, the collection volume was centered on the ninth and tenth myotomes. For all quantitative experiments, images of all samples were collected with identical laser power, camera gain, and exposure time.

### Longitudinal Imaging

To assess the *in vivo* persistence of mito-Keima fluorescence (Figure 6), WT embryos were injected with pTol2-*hsp70l:mito-Keima;cryaa:mCherry* to generate mosaic larvae. Larvae were fasted from 4.5 dpf to 6.5 dpf to induce a singular pulse of mitophagic flux. Larvae were heat shocked as described above in Fasting and Feeding Protocols to induce a singular pulse of mito-Keima protein expression. We then identified two larvae with suitable Keima expression patterns to allow for tracking of an individual muscle cell day after day. Each day the larvae were anesthetized with tricane, immobilized in 0.8% low melt-point agarose on glass bottom dishes for imaging, confocal imaging was performed using consistent laser power and exposure times, the larvae were recovered from the agar, and then returned to E3 medium and fed to prevent additional mitophagy.

### Image Analysis

All mito-Keima image analysis was conducting using Imaris (Bitplane) software with either FIJI (ImageJ) or MATLAB (MathWorks) plugins to enable image mathematics. Raw images from Nikon Elements (.nd2) were converted to Imaris (.ims) file format using Imaris Image Converter (Bitplane) and imported to Imaris. A surface mask was drawn manually using the Surfaces function to segment muscle tissue away from skin. A ratiometric image (FIJI or MATLAB) was generated by changing the data type to 32 Bit Float and applying the following equation: (Keima561 channel) / (Keima440 channel + 150). The 150 addition ensures that areas of low signal from both Keima561 and Keima440 were excluded. The ratiometric image normalizes for the decrease in signal intensity as imaging depth into the tissue increases and for small intensity differences between samples. Then, an isosurfaces thresholding algorithm was applied to the ratio channel, with 0.15 μm surface grain (smoothing) to eliminate noise, using a threshold value determined for each experiment from images of control fish. A size cutoff removed objects smaller than 2 voxels to eliminate additional noise. Obviously spurious surfaces were removed manually. Rarely, individual slices were removed from z-stack images to account for fish movement, which caused edge artifacts in some ratio images. Mitolysosome density was determined by dividing the number of ratio surface objects by the total tissue surface volume. Mitophagy index was determined by dividing the sum of all ratio surface object volumes by the total tissue surface volume. For mito-GR images, a deconvolution algorithm (Imaris) to adjust for the behavior of light of differing emission wavelengths (green and red) was used reduce edge artifacts. Object tracking in time-lapse microscopy (Figure 4 and Movie 2) was performed using Imaris software (Bitplane). Mitophagy isosurface masks were generated as described above. Mitochondria surfaces were generated using a 0.325 μm surface grain (smoothing), background subtraction (diameter largest sphere = 1.219 μm), a manually determined florescence intensity threshold, and a size cutoff eliminating objects smaller than 15 voxels. Object tracking was performed on ratio surface objects, using autoregressive motion, a maximum distance of 1.003 μm, a maximum gap size of 3 (without gap filling), and a track duration minimum of 5.534 s. We manually assessed each newly formed mitophagy isosurface object, defined as a Ratio^high^ tracked object that was not present at time zero; other mitophagy objects were considered stable. Using a combination of microscopy views, new objects were manually classified as new mitophagy, split mitolysosomes, or from outside. For lysosome size analysis, LysoTracker Green z-stack images were acquired covering 8.8 μm depth within the skeletal muscle of the fish and analyzed using FIJI. First, background subtraction using 50 μm rolling ball was performed, then a maximum intensity projection was generated, the area of muscle was isolated manually by drawing a gate, a threshold was automatically applied using the Triangle setting, touching objects were split using the watershed function, and objects were counted using the analyze particles function, cutting off objects smaller than 2 pixels. The number and area of lysosomes were measured using the maximum intensity projection image, and the area of the drawn gate served as total tissue area. For object-based colocalization experiments (Figure 5B-F, I), Keima was excited using 488 nm and 561 nm lasers using a microscope equipped with a 640 nm laser needed to excite LysoTracker Far Red (this microscope does not have a 440 nm laser). 488 nm exposure excites both acidic and neutral/basic mitochondria, whereas 561 nm excitation heavily favors excitation of acidic mitochondria. Thus, healthy mitochondria were segmented using an image generated using the following equation: (Keima488 channel) / (Keima561 channel + 1). One myotome was manually segmented, and all objects were counted within. Mitolysosomes were segmented using an image generated using the following equation (Keima561 channel) / (Keima488 channel + 150). EGFP-Lc3^+^ and LysoTracker Deep Red^+^ objects were segmented as described above. Object-based colocalization was performed using distance transformations from each object category, and objects near the edges of the somite or image border were excluded. Lc3^+^ objects were considered autolysosomes if there was a LysoTracker Deep Red object within 1 nm of it (filtered by Signal Intensity > 1 in the LysoTracker Deep Red objects distance transformation channel). If not, they were considered autophagosomes. An autophagosome or autolysosome was considered “in contact” with a healthy mitochondrion if there was a Keima488/561 object within 0 nm of it (filtered by Signal Intensity > 0 in the Keima488/561 objects distance transformation channel). A mitolysosome was considered near an Lc3^+^ object if it was within 0.5 μm (filtered by Signal Intensity > 500 in the Lc3^+^ objects distance transformation channel). For mito-Keima persistence experiments, individual cells were cropped and a Spot detection algorithm was applied to count Keima561^high^ puncta and track their decrease over time as Keima protein decayed.

### Morpholino Injections

Morpholinos (GeneTools, LLC) were reconstituted to 1 mM stock solutions in water. The injection mixes were prepared at 0.9 mM (*atg5*), 0.4 mM (vhl), or 0.2 mM (*pink1*), and 2 nL was injected into the yolk of 1-cell stage embryos. The sequences are listed in the Reagents and Resources Table.

### CRISPR

All CRISPR genome editing was performed using a variation of the high-efficiency protocol described (Burger *et al*, 2016). Guide RNA sequences were designed using CHOPCHOP (Labun *et al*, 2019) and chosen to target a 5’ exonic region with the potential to introduce STOP codons in the event of frameshift. Alt-R crRNA (IDT) constructs were reconstituted and annealed with Alt-R tracrRNA (IDT) to form gRNA duplexes according to the manufacturer’s instructions. Injection mixes were prepared by mixing 1 μL the gRNA duplex, 1 μL Duplex Buffer (IDT), and 1 μL ENGEN Spy Cas9 NLS (NEB) and allowing the mix to incubate at room temperature for at least 10 min. Approximately 2 nL of injection mix was injected into both the cell and yolk of 1-cell stage embryos. Efficient editing was confirmed by PCR and Sanger sequencing. For experiments using F0 CRISPR editing, the Sanger sequencing trace of each imaged larva was used to determine the extent of editing in a blinded manner. Editing was considered sufficient if trace files showed a mixture of multiple traces emerging near the PAM site. Images from larvae found to have low or no editing were removed from analysis (Supplemental Figure S5). To isolate the *bnip3* mutants, F0 parents were outcrossed and PCR and Sanger Sequencing was performed on gDNA isolated from pooled larvae and TOPO cloned into pCR4-TOPO TA vector (Invitrogen). Sequences were analyzed for the presence of alleles leading to premature stop codons and clutches containing such alleles were raised to adulthood and outcrossed. Adult F2 fish were tail clipped and subjected to similar TOPO sequencing to identify heterozygous founders.

### Genotyping

DNA from individual larvae was extracted using proteinase K (Invitrogen) and amplified by PCR with the primer pairs (IDT) listed in the Key Resources Table. The 5’ primer of the *pink1^gi2^* PCR genotyping primer pair was synthesized with a 5’ 6-FAM (Fluorescein) fluorescent dye attachment necessary for DNA Fragment Analysis, in which small, fluorescently-labeled DNA fragments are electrophoretically separated such that precise changes in size (down to 1 bp) can be detected. The *pink1^gi2^* (Zhang *et al*, 2017) and *bnip3* mutant fish were identified by the presence of a small deletion, which was detected using DNA Fragment Analysis (MGH CCIB DNA Core). Efficient editing in F0 high-efficiency CRISPR experiments was confirmed by Sanger sequencing, as described in the CRISPR section.

### Statistical Analyses

All graphs and statistical analyses were generated using Prism (GraphPad). All bar graphs display the mean + SEM unless otherwise stated in the figure legend. For comparisons between only two conditions, standard, unpaired, two-tailed Student’s t-tests were employed. For comparisons within groups with 4 conditions, two-way ANOVA with Holm-Sidak’s multiple comparisons test were employed. *P* values or an asterisk indicating significant differences as defined in the figure legends were included in each figure containing statistical analyses. Supplemental Figures S4, S6-S8 were included for specific quantitative experiments to demonstrate reproducibility and provide full statistical analyses.

## ACKNOWLEDGEMENTS

Research was supported by NIH F32AA025271 and the American Liver Foundation (P.J.W.), NIH R37 NS083524 to J.W.H., and NIH R01DK090311, R01DK105198, R24OD017870 to W.G.. W.G. is a Pew Scholar in the Biomedical Sciences. We thank Tom Schwarz for helpful discussions. We thank Nadine Budrow for maintaining our zebrafish colonies. We thank Phil Elks for sharing *DA-hif* plasmids, Donghun Shin for sharing the p5E-2.8fabp10a plasmid, and Daniel Hesselson for sharing the *pink1*^gi2^ line. We thank the Harvard Medical School MicRoN team for access to microscopes and imaging expertise.

## AUTHOR CONTRIBUTIONS

Conceptualization, P.J.W., J-M.H., J.W.H., and W.G.; Methodology, P.J.W., A.S., and J-M.H.; Investigation, P.J.W., E.D.Q., and K.A.L.; Writing – Original Draft, P.J.W. and W.G.; Writing – Review and Editing, P.J.W., A.S., J-M.H., E.D.Q., J.W.H. and W.G.; Supervision, J.W.H. and W.G.; Funding Acquisition, P.J.W., J.W.H., and W.G.

## REFERENCES

Abeliovich H, Zarei M, Rigbolt KTG, Youle RJ & Dengjel J (2013) Involvement of mitochondrial dynamics in the segregation of mitochondrial matrix proteins during stationary phase mitophagy. Nat Commun 4: 2789

Abudu YP, Pankiv S, Mathai BJ, Håkon Lystad A, Bindesbøll C, Brenne HB, Yoke Wui Ng M, Thiede B, Yamamoto A, Mutugi Nthiga T, et al (2019) NIPSNAP1 and NIPSNAP2 Act as “Eat Me” Signals for Mitophagy. Dev Cell 49: 509–525.e12

Anding AL & Baehrecke EH (2017) Cleaning house: Selective autophagy of organelles. Dev Cell 41: 10–22

Berger J & Currie PD (2013) 503Unc, a Small and Muscle-Specific Zebrafish Promoter. Genesis 51: 443–447

Bill BR, Petzold AM, Clark KJ, Schimmenti LA & Ekker SC (2009) A primer for morpholino use in zebrafish. Zebrafish 6: 69–77

Birgisdottir A, Lamark T & Johansen T (2013) The LIR motif – crucial for selective autophagy. J Cell Sci 126: 3237–3247

Burger A, Lindsay H, Felker A, Hess C, Anders C, Chiavacci E, Zaugg J, Weber LM, Catena R, Jinek M, et al (2016) Maximizing mutagenesis with solubilized CRISPR-Cas9 ribonucleoprotein complexes. Development 143: 2025–2037

Burman JL, Pickles S, Wang C, Sekine S, Vargas JNS, Zhang Z, Youle AM, Nezich CL, Wu X, Hammer JA, et al (2017) Mitochondrial fission facilitates the selective mitophagy of protein aggregates. J Cell Biol 216: 3231–3247

Cadete VJJ, Deschênes S, Cuillerier A, Brisebois F, Sugiura A, Vincent A, Turnbull D, Picard M, McBride HM & Burelle Y (2016) Formation of mitochondrial-derived vesicles is an active and physiologically relevant mitochondrial quality control process in the cardiac system. J Physiol 594: 5343–5362

Chen G, Han Z, Feng D, Chen Y, Chen L, Wu H, Huang L, Zhou C, Cai X, Fu C, et al (2014) A regulatory signaling loop comprising the PGAM5 phosphatase and CK2 controls receptor-mediated mitophagy. Mol Cell 54: 362–77

Clark IE, Dodson MW, Jiang C, Cao JH, Huh JR, Seol JH, Yoo SJ, Hay BA & Guo M (2006) Drosophila pink1 is required for mitochondrial function and interacts genetically with parkin. Nature 441: 1162–1166

Cox AG & Goessling W (2015) The lure of zebrafish in liver research: Regulation of hepatic growth in development and regeneration. Curr Opin Genet Dev 32: 153–161

DiNapoli SE, Martinez-McFaline R, Gribbin CK, Wrighton PJ, Balgobin CA, Nelson I, Leonard A, Maskin CR, Shwartz A, Quenzer ED, et al (2020) Synthetic CRISPR/Cas9 reagents facilitate genome editing and homology directed repair. Nucleic Acids Res Epub

Elks PM, Brizee S, van der Vaart M, Walmsley SR, van Eeden FJ, Renshaw SA & Meijer AH (2013) Hypoxia Inducible Factor Signaling Modulates Susceptibility to Mycobacterial Infection via a Nitric Oxide Dependent Mechanism. PLoS Pathog 9: e1003789

Elks PM, Van Eeden FJ, Dixon G, Wang X, Reyes-Aldasoro CC, Ingham PW, Whyte MKB, Walmsley SR & Renshaw SA (2011) Activation of hypoxia-inducible factor-1α (hif-1α) delays inflammation resolution by reducing neutrophil apoptosis and reverse migration in a zebrafish inflammation model. Blood 118: 712–722

Evans CS & Holzbaur EL (2020) Degradation of engulfed mitochondria is rate-limiting in Optineurin-mediated mitophagy in neurons. Elife 9: e50260

Ferdinand P & Roffe C (2016) Hypoxia after stroke: A review of experimental and clinical evidence. Exp Transl Stroke Med 8: 1–8

Friedman JR & Nunnari J (2014) Mitochondrial form and function. Nature 505: 335–343

Gasser T (2009) Mendelian forms of Parkinson’s disease. Biochim Biophys Acta - Mol Basis Dis 1792: 587–596

Le Guerroué F, Eck F, Jung J, Starzetz T, Mittelbronn M, Kaulich M & Behrends C (2017) Autophagosomal Content Profiling Reveals an LC3C-Dependent Piecemeal Mitophagy Pathway. Mol Cell 68: 786–796.e6

Hamacher-Brady A, Brady NR, Logue SE, Sayen MR, Jinno M, Kirshenbaum LA, Gottlieb RA & Gustafsson ÅB (2007) Response to myocardial ischemia/reperfusion injury involves Bnip3 and autophagy. Cell Death Differ 14: 146–157

Hanna RA, Quinsay MN, Orogo AM, Giang K, Rikka S & Gustafsson ÅB (2012) Microtubule-associated protein 1 light chain 3 (LC3) interacts with Bnip3 protein to selectively remove endoplasmic reticulum and mitochondria via autophagy. J Biol Chem

Harper JW, Ordureau A & Heo JM (2018) Building and decoding ubiquitin chains for mitophagy. Nat Rev Mol Cell Biol 19: 93–108

Harris JM, Esain V, Frechette GM, Harris LJ, Cox AG, Cortes M, Garnaas MK, Carroll KJ, Cutting CC, Khan T, et al (2013) Glucose metabolism impacts the spatiotemporal onset and magnitude of HSC induction in vivo. Blood 121: 2483–2493

He C, Bartholomew CR, Zhou W & Klionsky DJ (2009) Assaying autophagic activity in transgenic GFP-Lc3 and GFP-Gabarap zebrafish embryos. Autophagy 5: 520–526

Her GM, Chiang C-C, Chen W-Y & Wu J-L (2003) In vivo studies of liver-type fatty acid binding protein (L-FABP) gene expression in liver of transgenic zebrafish (Danio rerio). FEBS Lett 538: 125–133

Iliopoulos O, Levy AP, Jiang C, Kaelin WG & Goldberg MA (1996) Negative regulation of hypoxia-inducible genes by the von Hippel-Lindau protein. Proc Natl Acad Sci 93: 10595–10599

Ivan M, Kondo K, Yang H, Kim W, Valiando J, Ohh M, Salic A, Asara JM, Lane WS & Kaelin J (2001) HIFα targeted for VHL-mediated destruction by proline hydroxylation: Implications for O2 sensing. Science 292: 464–468

Kaelin WG & McKnight SL (2013) Influence of metabolism on epigenetics and disease. Cell 153: 56–69

Katayama H, Hama H, Nagasawa K, Kurokawa H, Sugiyama M, Ando R, Funata M, Yoshida N, Homma M, Nishimura T, et al (2020) Visualizing and Modulating Mitophagy for Therapeutic Studies of Neurodegeneration. Cell 181: 1176–1187.e16

Katayama H, Kogure T, Mizushima N, Yoshimori T & Miyawaki A (2011) A sensitive and quantitative technique for detecting autophagic events based on lysosomal delivery. Chem Biol 18: 1042–1052

Katayama H, Yamamoto A, Mizushima N, Yoshimori T & Miyawaki A (2008) GFP-like proteins stably accumulate in lysosomes. Cell Struct Funct 33: 1–12

Kim YY, Um J, Yoon J, Kim H, Lee YJ, Jee HJ, Kim YM, Jang JS, Jang Y, Chung J, et al (2019) Assessment of mitophagy in mt-Keima Drosophila revealed an essential role of the PINK1-Parkin pathway in mitophagy induction in vivo. FASEB J 33: 9742–9751

Kolitsida P, Zhou J, Rackiewicz M, Nolic V, Dengjel J & Abeliovich H (2019) Phosphorylation of mitochondrial matrix proteins regulates their selective mitophagic degradation. Proc Natl Acad Sci 116: 20517–20527

Kwan KM, Fujimoto E, Grabher C, Mangum BD, Hardy ME, Campbell DS, Parant JM, Yost HJ, Kanki JP & Chien C Bin (2007) The Tol2kit: A multisite gateway-based construction Kit for Tol2 transposon transgenesis constructs. Dev Dyn 236: 3088–3099

Labun K, Montague TG, Krause M, Torres Cleuren YN, Tjeldnes H & Valen E (2019) CHOPCHOP v3: expanding the CRISPR web toolbox beyond genome editing. Nucleic Acids Res 47: W171–W174

Lee E, Koo Y, Ng A, Wei Y, Luby-Phelps K, Juraszek A, Xavier RJ, Cleaver O, Levine B & Amatruda JF (2014) Autophagy is essential for cardiac morphogenesis during vertebrate development. Autophagy 10: 572–587

Lee E, Wei Y, Zou Z, Tucker K, Rakheja D, Levine B & Amatruda JF (2016) Genetic inhibition of autophagy promotes p53 loss-of-heterozygosity and tumorigenesis. Oncotarget 7

Lee JJ, Sanchez-Martinez A, Zarate AM, Benincá C, Mayor U, Clague MJ & Whitworth AJ (2018) Basal mitophagy is widespread in Drosophila but minimally affected by loss of Pink1 or parkin. J Cell Biol 217: 1613–1622

Lee JW, Ko J, Ju C & Eltzschig HK (2019) Hypoxia signaling in human diseases and therapeutic targets. Exp Mol Med 51: 1–13

Leermakers PA & Gosker HR (2016) Skeletal muscle mitophagy in chronic disease: Implications for muscle oxidative capacity. Curr Opin Clin Nutr Metab Care 19: 427–433

Liu L, Feng D, Chen G, Chen M, Zheng Q, Song P, Ma Q, Zhu C, Wang R, Qi W, et al (2012) Mitochondrial outer-membrane protein FUNDC1 mediates hypoxia-induced mitophagy in mammalian cells. Nat Cell Biol 14: 177–185

Mammucari C, Milan G, Romanello V, Masiero E, Rudolf R, Del Piccolo P, Burden SJ, Di Lisi R, Sandri C, Zhao J, et al (2007) FoxO3 Controls Autophagy in Skeletal Muscle In Vivo. Cell Metab 6: 458–471

Mao K, Wang K, Zhao M, Xu T & Klionsky DJ (2011) Two MAPK-signaling pathways are required for mitophagy in Saccharomyces cerevisiae. J Cell Biol 193: 755–767

McWilliams TG, Prescott AR, Allen GFG, Tamjar J, Munson MJ, Thomson C, Muqit MMK & Ganley IG (2016) Mito-QC illuminates mitophagy and mitochondrial architecture in vivo. J Cell Biol 214: 333–345

McWilliams TG, Prescott AR, Montava-Garriga L, Ball G, Singh F, Barini E, Muqit MMK, Brooks SP & Ganley IG (2018) Basal Mitophagy Occurs Independently of PINK1 in Mouse Tissues of High Metabolic Demand. Cell Metab 27: 439–449.e5

Mosimann C, Kaufman CK, Li P, Pugach EK, Tamplin OJ & Zon LI (2011) Ubiquitous transgene expression and Cre-based recombination driven by the ubiquitin promoter in zebrafish. Development 138: 169–177

Mosimann C & Zon LI (2011) Advanced zebrafish transgenesis with Tol2 and application for Cre/lox recombination experiments. Methods Cell Biol 104: 173–194

Ordureau A, Paulo JA, Zhang J, An H, Swatek KN, Cannon JR, Wan Q, Komander D & Harper JW (2020) Global Landscape and Dynamics of Parkin and USP30-Dependent Ubiquitylomes in iNeurons during Mitophagic Signaling. Mol Cell 77: 1124–1142.e10

Palikaras K, Lionaki E & Tavernarakis N (2018) Mechanisms of mitophagy in cellular homeostasis, physiology and pathology. Nat Cell Biol 20: 1013–1022

Park J, Lee SB, Lee S, Kim Y, Song S, Kim S, Bae E, Kim J, Shong M, Kim JM, et al (2006) Mitochondrial dysfunction in Drosophila PINK1 mutants is complemented by parkin. Nature 441: 1157–1161

Pickles S, Vigié P & Youle RJ (2018) Mitophagy and Quality Control Mechanisms in Mitochondrial Maintenance. Curr Biol 28: R170–R185

Pickrell AM & Youle RJ (2015) The roles of PINK1, Parkin, and mitochondrial fidelity in parkinson’s disease. Neuron 85: 257–273

Porcelli AM, Ghelli A, Zanna C, Pinton P, Rizzuto R & Rugolo M (2005) pH difference across the outer mitochondrial membrane measured with a green fluorescent protein mutant. Biochem Biophys Res Commun 326: 799–804

Preidis GA, Kim KH & Moore DD (2017) Nutrient-sensing nuclear receptors PPAR? and FXR control liver energy balance. J Clin Invest 127: 1193–1201

Priyadarshini M, Tuimala J, Chen YC & Panula P (2013) A zebrafish model of PINK1 deficiency reveals key pathway dysfunction including HIF signaling. Neurobiol Dis 54: 127–138

Raefsky SM & Mattson MP (2017) Adaptive responses of neuronal mitochondria to bioenergetic challenges: Roles in neuroplasticity and disease resistance. Free Radic Biol Med 102: 203–216

Rodger CE, Mcwilliams TG, Ganley IG & Ganley IG (2018) Mammalian mitophagy – from in vitro molecules to in vivo models. FEBS J 285: 1185–1202

Rodríguez-Nuevo A, Díaz-Ramos A, Noguera E, Díaz-Sáez F, Duran X, Muñoz JP, Romero M, Plana N, Sebastián D, Tezze C, et al (2018) Mitochondrial DNA and TLR9 drive muscle inflammation upon Opa1 deficiency. EMBO J 37: 1–18

Rojansky R, Cha MY & Chan DC (2016) Elimination of paternal mitochondria in mouse embryos occurs through autophagic degradation dependent on PARKIN and MUL1. Elife 5: e17896

Romanello V & Sandri M (2013) Mitochondrial biogenesis and fragmentation as regulators of protein degradation in striated muscles. J Mol Cell Cardiol 55: 64–72

Ronan B, Flamand O, Vescovi L, Dureuil C, Durand L, Fassy F, Bachelot MF, Lamberton A, Mathieu M, Bertrand T, et al (2014) A highly potent and selective Vps34 inhibitor alters vesicle trafficking and autophagy. Nat Chem Biol 10: 1013–1019

Sandri M (2013) Protein breakdown in muscle wasting: Role of autophagy-lysosome and ubiquitin-proteasome. Int J Biochem Cell Biol 45: 2121–2129

Sebastián D & Zorzano A (2020) Self-Eating for Muscle Fitness: Autophagy in the Control of Energy Metabolism. Dev Cell 54: 268–281

Settembre C, Fraldi A, Medina DL & Ballabio A (2013) Signals from the lysosome: A control centre for cellular clearance and energy metabolism. Nat Rev Mol Cell Biol 14: 283–296

Shi RY, Zhu SH, Li V, Gibson SB, Xu XS & Kong JM (2014) BNIP3 interacting with LC3 triggers excessive mitophagy in delayed neuronal death in stroke. CNS Neurosci Ther 20: 1045–1055

Smirnova E, Shurland DL, Ryazantsev SN & Van Der Bliek AM (1998) A human dynamin-related protein controls the distribution of mitochondria. J Cell Biol 143: 351–358

Sugiura A, McLelland G, Fon EA & McBride HM (2014) A new pathway for mitochondrial quality control: mitochondrial-derived vesicles. EMBO J 33: 2142–2156

Sun N, Yun J, Liu J, Malide D, Liu C, Rovira II, Holmström KM, Fergusson MM, Yoo YH, Combs CA, et al (2015) Measuring In Vivo Mitophagy. Mol Cell 60: 685–696

Vargas JNS, Wang C, Bunker E, Hao L, Maric D, Schiavo G, Randow F & Youle RJ (2019) Spatiotemporal Control of ULK1 Activation by NDP52 and TBK1 during Selective Autophagy. Mol Cell 74: 347–362.e6

Williams JA, Ni HM, Haynes A, Manley S, Li Y, Jaeschke H & Ding WX (2015) Chronic deletion and acute knockdown of Parkin have differential responses to acetaminophen-induced mitophagy and liver injury in mice. J Biol Chem 290: 10934–10946

Wilson C (2012) Aspects of larval rearing. ILAR J 53: 169–178

Wrighton PJ, Oderberg IM & Goessling W (2019) There Is Something Fishy About Liver Cancer: Zebrafish Models of Hepatocellular Carcinoma. Cell Mol Gastroenterol Hepatol 8: 347–363

Yamashita SI, Jin X, Furukawa K, Hamasaki M, Nezu A, Otera H, Saigusa T, Yoshimori T, Sakai Y, Mihara K, et al (2016) Mitochondrial division occurs concurrently with autophagosome formation but independently of Drp1 during mitophagy. J Cell Biol 215: 649–665

Yang W, Liu Y, Tu Z, Xiao C, Yan S, Ma X, Guo X, Chen X, Yin P, Yang Z, et al (2019) CRISPR/Cas9-mediated PINK1 deletion leads to neurodegeneration in rhesus monkeys. Cell Res 29: 334–336

Yang Y, Gehrke S, Imai Y, Huang Z, Ouyang Y, Wang JW, Yang L, Beal MF, Vogel H & Lu B (2006) Mitochondrial pathology and muscle and dopaminergic neuron degeneration caused by inactivation of Drosophila Pink1 is rescued by Parkin. Proc Natl Acad Sci 103: 10793–10798

Yoo S-M & Jung Y-K (2018) A Molecular Approach to Mitophagy and Mitochondrial Dynamics. Mol Cells 41: 18–26

Youle RJ (2019) Mitochondria—Striking a balance between host and endosymbiont. Science 365

Yuan Y, Zheng Y, Zhang X, Chen Y, Wu X, Wu J, Shen Z, Jiang L, Wang L, Yang W, et al (2017) BNIP3L/NIX-mediated mitophagy protects against ischemic brain injury independent of PARK2. Autophagy 13: 1754–1766

Zhang H, Bosch-Marce M, Shimoda LA, Yee ST, Jin HB, Wesley JB, Gonzalez FJ & Semenza GL (2008) Mitochondrial autophagy is an HIF-1-dependent adaptive metabolic response to hypoxia. J Biol Chem 283: 10892–10903

Zhang T, Xue L, Li L, Tang C, Wan Z, Wang R, Tan J, Tan Y, Han H, Tian R, et al (2016a) BNIP3 protein suppresses PINK1 kinase proteolytic cleavage to promote mitophagy. J Biol Chem 291: 21616–21629

Zhang W, Ren H, Xu C, Zhu C, Wu H, Liu D, Wang J, Liu L, Li W, Ma Q, et al (2016b) Hypoxic mitophagy regulates mitochondrial quality and platelet activation and determines severity of I/R heart injury. Elife 5: e21407

Zhang Y, Nguyen DT, Olzomer EM, Poon GP, Cole NJ, Puvanendran A, Phillips BR & Hesselson D (2017) Rescue of Pink1 Deficiency by Stress-Dependent Activation of Autophagy. Cell Chem Biol 24: 471–480.e4

Zhou H, Zhu P, Guo J, Hu N, Wang S, Li D, Hu S, Ren J, Cao F & Chen Y (2017) Ripk3 induces mitochondrial apoptosis via inhibition of FUNDC1 mitophagy in cardiac IR injury. Redox Biol 13: 498–507

Zhou Y, Long Q, Wu H, Li W, Qi J, Wu Y, Xiang G, Tang H, Yang L, Chen K, et al (2019) Topology-dependent, bifurcated mitochondrial quality control under starvation. Autophagy Jul: 1–13

