## Supplemental Information for "Quantitative intravital imaging reveals *in vivo* dynamics of physiological-stress induced mitophagy"

**Resources Table**

| REAGENT or RESOURCE | SOURCE | IDENTIFIER |
|---|---|---|
| **Experimental Models: Organisms/Strains** | | |
| Zebrafish: *Tg(CMV:EGFP-map1lc3b)zf155* | (He *et al*, 2009) | ZFIN ID: ZDB-GENO-091029-2 |
| Zebrafish: *Tg(ubi:mito-Keima)* | This paper. | N/A |
| Zebrafish: *Tg(ubi:mito-GR)* | This paper. | N/A |
| Zebrafish: *Tg(fabp10a:mito-GR)* | This paper. | N/A |
| **Recombinant DNA** | | |
| Plasmid: pTol2-*ubi:mito-Keima,cryaa:Cerulean* | This paper. | N/A |
| Plasmid: pTol2-*ubi:mito-GR* | This paper. | N/A |
| Plasmid: pTol2--*2.8fabp10a:mito-GR* | This paper. | N/A |
| Plasmid: pHAGE mt-Keima IRES Puro | (Katayama *et al*, 2011) | N/A |
| Plasmid: pENTR-mito-Keima | This paper. | N/A |
| Plasmid: pCLBW cox8 EGFP mCherry | (Rojansky *et al*, 2016) | Addgene Plasmid #78520 |
| Plasmid: pME-mito-GR | This paper. | N/A |
| Plasmid: Gateway pDONR221 | ThermoFisher | Cat#: 12536017 |
| Plasmid: pENTR5'_ubi | (Mosimann *et al*, 2011) | Addgene Plasmid #27320 |
| Plasmid: p5E-2.8fabp10a | (Choi *et al*, 2014) | Gift from Donghun Shin |

| | | |
|---|---|---|
| Plasmid: p3E-2A-pCS2MCS-pA | (Kwan *et al*, 2007) | Tol2 kit v2.0 plasmid #633 |
| Plasmid: pDEST-tol2-cryaaCerulean | (Kwan *et al*, 2007) | N/A |
| Plasmid: pDestTol2pA2 | (Kwan *et al*, 2007) | Tol2 kit v1.0 plasmid #394 |
| Plasmid: pCS2_da-hif1ab | (Elks *et al*, 2011) | Gift from Phil Elks. |
| Plasmid: p5E_unc503 | (Berger & Currie, 2013) | Addgene Plasmid #64020 |
| Plasmid: p3E-2A-nlsEGFPpA | (Kwan *et al*, 2007) | Tol2 kit v2.0 plasmid #459 |
| Plasmid: p3E-MCS | (Don *et al*, 2017) | Addgene Plasmid #75174 |
| Plasmid: pDESTtol2pACrymCherry | (Berger & Currie, 2013) | Addgene Plasmid #64023 |
| Plasmid: p5E-hsp70l | (Kwan *et al*, 2007) | Tol2 kit v1.0 plasmid #222 |
| Plasmid: pENTR-DA-hif1ab(noStop) | This paper. | N/A |
| Plasmid: pTol2--*503unc45b:DA-hif1ab-2A-nlsEGFP,cryaa:mCherry* | This paper. | N/A |
| Plasmid: pENTR-DN-Drp1(noStop) | This paper. | N/A |
| Plasmid: pTol2-*hsp70l:DN-Drp1-2A-nlsEGFP,cryaa:mCherry* | This paper. | N/A |
| Plasmid: pTol2-*hsp70l:mito-Keima,cryaa:mCherry* | This paper. | N/A |

| | | |
|---|---|---|
| Plasmid: pCS2FA-transposase | (Kwan *et al*, 2007) | Tol2 kit v1.0 plasmid #396 |
| **Chemicals, Peptides, and Recombinant Proteins** | | |
| Dimethyl Sulfoxide (DMSO) | Sigma Aldrich | |
| DMOG | Tocris | Cat#: 4408 |
| FG-4592 | Cayman Chemical | Cat#: 15294 |
| MitoTracker Green FM | Cell Signaling Technology | Cat#: 9074S |
| LysoTracker Green DND-26 | Cell Signaling Technology | Cat#: 8783S |
| LysoTracker Deep Red | Thermo Fisher Scientific | Cat#: L12492 |
| Tricane-S | Western Chemical (Thermo Fisher Scientific) | Cat#: NC0872873 |
| Low Melting Point Agarose | Invitrogen (Thermo Fisher Scientific) | Cat#: 16-520-100 |
| Engen Spy Cas9 NLS | New England Biolabs | Cat#: M0646M |
| LR Clonase II Plus enzyme | Invitrogen | Cat#: 12538120 |
| Gateway BP Clonase II Enzyme mix | Invitrogen | Cat#: 11789020 |
| **Oligonucleotides** | | |
| Morpholino: Standard Control Oligo CCTCTTACCTCAGTTACAATTTATA | GeneTools, LLC | Standard Negative Control Oligo |

| | | |
|---|---|---|
| Morpholino: *atg5*<br>CACATCCTTGTCATCTGCCATTATC | (Lee *et al*, 2014, 2016) | GeneTools, LLC |
| Morpholino: *pink1*<br>TCACAACCTACCCGTTCAAAGTCAG | (Priyadarshini *et al*, 2013) | GeneTools, LLC |
| Morpholino: *vhl*<br>GGCATCGTCAAAGACAGGACAGTTC | (Harris *et al*, 2013) | GeneTools, LLC |
| Alt-R CRISPR-Cas9 crRNA: *pink1*<br>TCTTCAGGTTGTCTGTCAGC | This paper. | IDT |
| Alt-R CRISPR-Cas9 crRNA: *bnip3*<br>ATGCTCAGCACGAGTCGGGT | This paper. | IDT |
| Alt-R CRISPR-Cas9 crRNA: *bnip3la* (*nix*)<br>CTGCTGGGGAACGGGAACCA | This paper. | IDT |
| Alt-R CRISPR-Cas9 crRNA: *fundc1*<br>CAATGGTGGAGTCGAGTATT | This paper. | IDT |
| Alt-R CRISPR-Cas9 crRNA: *prkn*<br>GTGCGGTTTAATTCCAGCCA | This paper. | IDT |
| Alt-R CRISPR-Cas9 crRNA: Non-targeting Control<br>TGCTGATGTCTTGGTTGAAT | IDT | N/A |
| Alt-R CRISPR-Cas9 tracrRNA | IDT | Cat.#: 1072532 |
| PCR Genotyping Primer Pair: *pink1 (gi2)*<br>5': /6-FAM/CGCTCACAGAGACCTCAAAT<br>3': GGGAAGGGTAAAGGAGTAGA | This paper. | IDT |
| PCR Genotyping Primer Pair: *bnip3*<br>5': GTTCTTGGGTGGAGCTGCAT | This paper. | IDT |

| | | |
|---|---|---|
| 3': TGTCACATGGTAGGCTTCCT | | |
| PCR Genotyping Primer Pair: *bnip3la* (*nix*)<br><br>5': CTGGGTGGAGCTGGAGATGAAT<br><br>3': CTGTTGCAGGACGAGCTGCT | This paper. | IDT |
| PCR Genotyping Primer Pair: *fundc1*<br><br>5': GCAGAGAGTGAAGATGAATTG<br><br>3': AGCTTTCCAACTCTCTGAAA | This paper. | IDT |
| PCR Genotyping Primer Pair: *prkn*<br><br>5': GCTGTCTGAGGAGGTTTACC<br><br>3': CCCTGTGTCAGCATCAGAATTC | This paper. | IDT |

### Supplemental Figures

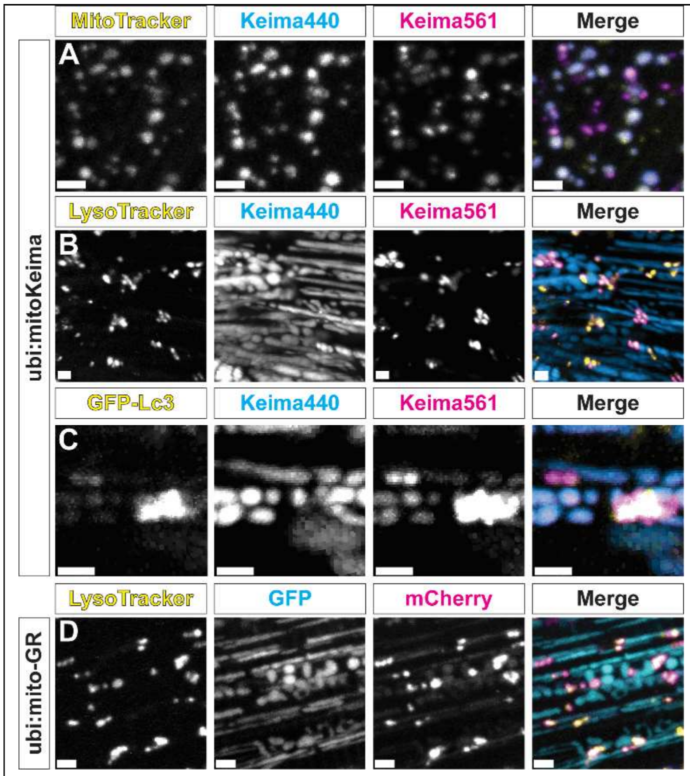


**Figure S1.** Reporter zebrafish faithfully differentiate healthy mitochondria and those undergoing mitophagy. (A) Maximum intensity projection (MIP) confocal images of skin cells of ubi:mito-Keima zebrafish larva treated with

MitoTracker Green, showing colocalization of MitoTracker puncta with both Keima440$^{high}$ puncta and Keima561$^{high}$ puncta. (B) MIP confocal images of skeletal muscle cells in ubi:mito-Keima zebrafish larva treated with LysoTracker Green, showing colocalization between the Keima561$^{high}$ puncta and LysoTracker puncta. (C) MIP confocal images of a ubi:mito-Keima;CMV:GFP-Lc3 zebrafish larva, demonstrating colocalization between Lc3$^{+}$ puncta and Keima561$^{high}$ puncta. (D) MIP confocal images of a ubi:mito-GR zebrafish larva treated with LysoTracker Deep Red, demonstrating colocalization between LysoTracker and the mCherry$^{+}$;EGFP$^{-}$ puncta. Scale bars, 3 μm.

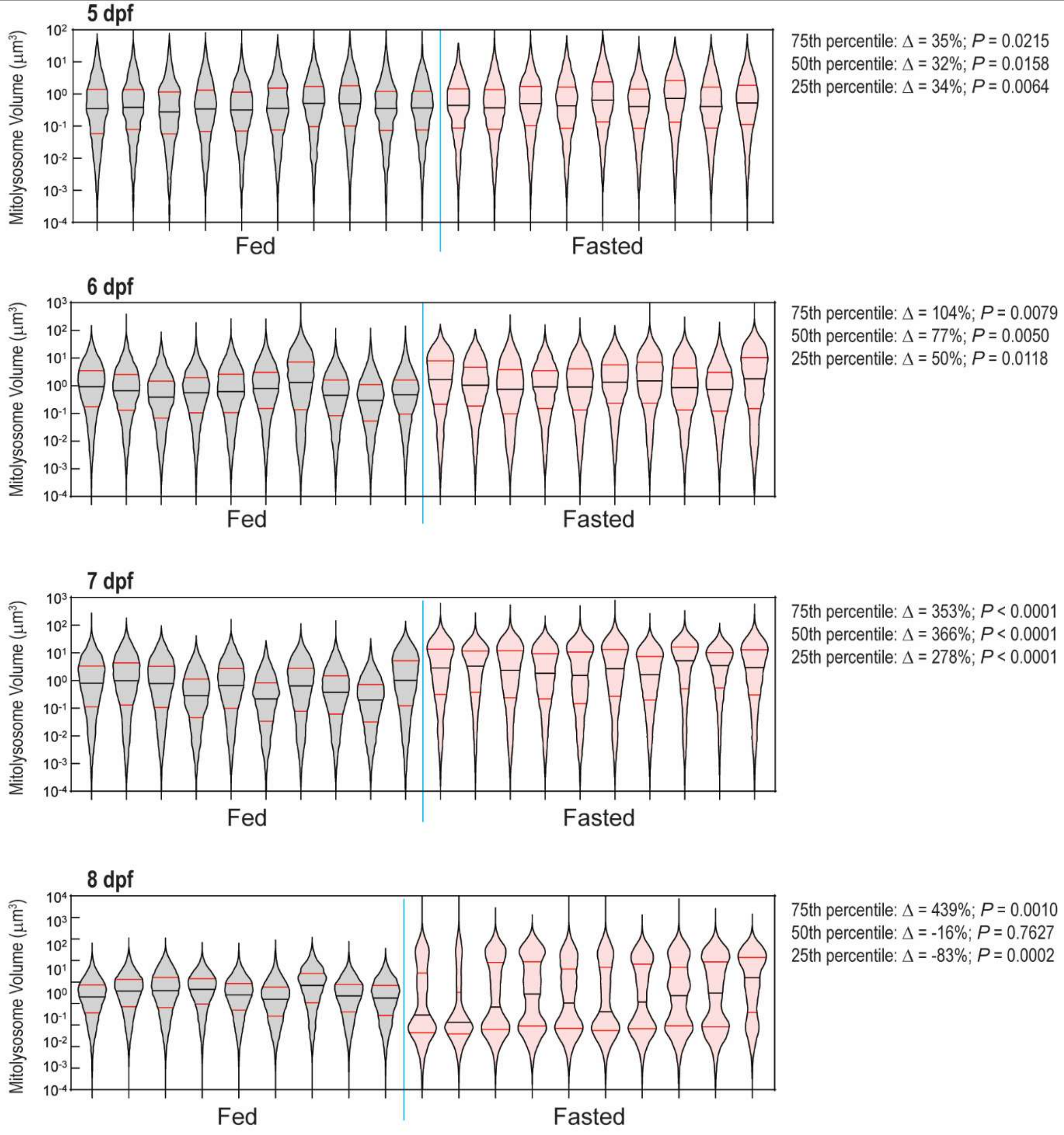


**Figure S2.** Volumetric distribution analyses reveal early and late changes in mitolysosome dynamics during fasting.

Violin plots of the mitolysosomes volumes (μm$^3$) of individual fed and fasted fish depicted in Figure 3. A significant increase in the volume of the 25th, 50th, and 75th percentile mitolysosome is evident at 5 dpf and continues through 7 dpf. At 8 dpf, the fasted cohort displays a bimodal mitolysosome volume distribution indicative of an increase in both large and small mitolysosomes. Data shown are violin distribution plots. Black line is 50th percentile. Red lines are 25th and 75th percentiles. *P* is unpaired, two-tailed Student's t test comparing the indicated percentile in fasted and fed cohorts. Δ is the percentage change of the indicated percentile in each Fasted compared to Fed cohort.

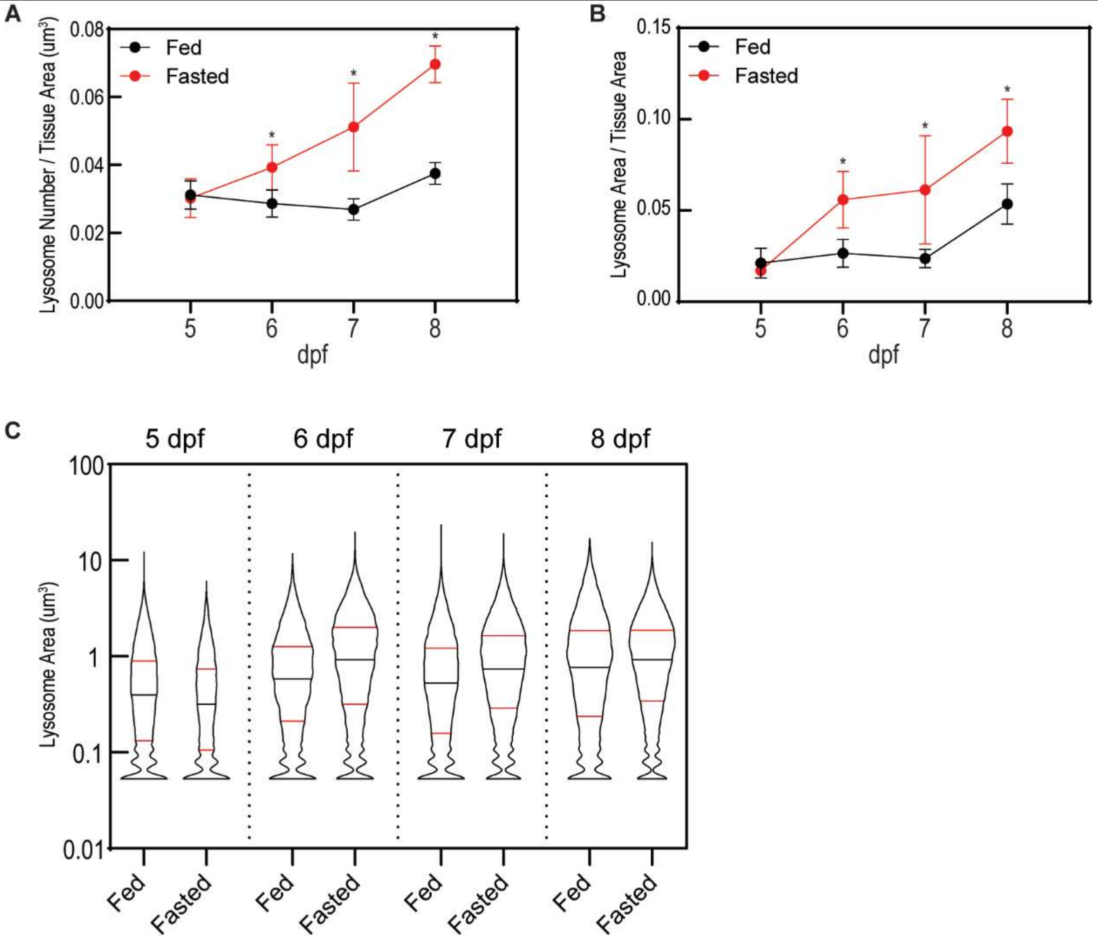


**Figure S3.** Lysosomes increase in size and number during fasting, but very small lysosomes are always present. (A) Tracking the density of lysosomes in skeletal muscle in fed and fasted fish from 5 to 8 dpf. Data are the mean lysosome density calculated from at least 7 larvae. Bars show standard deviation, and * indicates *P*<0.001 (unpaired, two-tailed Student's t test) between the fed and fasted cohorts within each time point. (B) Tracking the total area of all lysosomes within the muscle. Data are the mean lysosome area per total tissue area calculated from at least 7 larvae—same larvae as (A). Bars show standard deviation, and * indicates *P*<0.002 (unpaired, two-tailed Student's t test) between the fed and fasted cohorts within each time point. (*C*) Violin plots demonstrating the area distribution of lysosomes in fed and fasted muscle from 5 to 8 dpf.

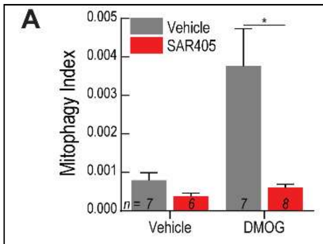


**Same as Fig. 7A**

| *Holm-Sidak's multiple comparisons test* | *Predicted (LS) mean diff.* | *Significant?* | *Summary* | *Adjusted P Value* |
|---|---|---|---|---|
| Vehicle:Vehicle vs. Vehicle:SAR405 | 0.0004188 | No | ns | 0.9232 |
| Vehicle:Vehicle vs. DMOG:Vehicle | -0.002963 | Yes | ** | 0.0013 |
| Vehicle:Vehicle vs. DMOG:SAR405 | 0.0001934 | No | ns | 0.9401 |
| Vehicle:SAR405 vs. DMOG:Vehicle | -0.003382 | Yes | *** | 0.0007 |
| Vehicle:SAR405 vs. DMOG:SAR405 | -0.0002254 | No | ns | 0.9401 |
| DMOG:Vehicle vs. DMOG:SAR405 | 0.003156 | Yes | *** | 0.0007 |

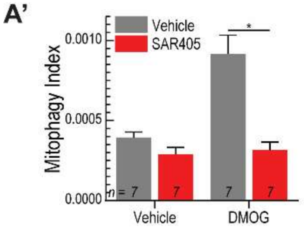


**Repeat of Fig. 7A**

| *Holm-Sidak's multiple comparisons test* | *Predicted (LS) mean diff.* | *Significant?* | *Summary* | *Adjusted P Value* |
|---|---|---|---|---|
| Vehicle:Vehicle vs. Vehicle:SAR405 | 0.0001054 | No | ns | 0.6545 |
| Vehicle:Vehicle vs. DMOG:Vehicle | -0.0005224 | Yes | **** | <0.0001 |
| Vehicle:Vehicle vs. DMOG:SAR405 | 7.899e-005 | No | ns | 0.6789 |
| Vehicle:SAR405 vs. DMOG:Vehicle | -0.0006275 | Yes | **** | <0.0001 |
| Vehicle:SAR405 vs. DMOG:SAR405 | -2.640e-005 | No | ns | 0.7923 |
| DMOG:Vehicle vs. DMOG:SAR405 | 0.0006014 | Yes | **** | <0.0001 |

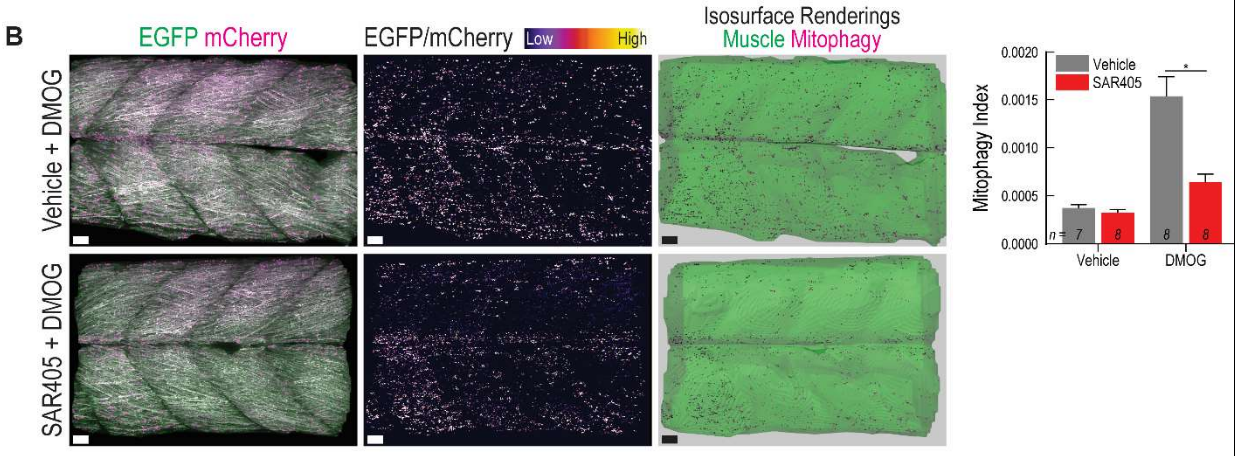


| *Holm-Sidak's multiple comparisons test* | *Predicted (LS) mean diff.* | *Significant?* | *Summary* | *Adjusted P Value* |
|---|---|---|---|---|
| Vehicle:Vehicle vs. Vehicle:SAR405 | 4.804e-005 | No | ns | 0.7830 |
| Vehicle:Vehicle vs. DMOG:Vehicle | -0.001161 | Yes | **** | <0.0001 |
| Vehicle:Vehicle vs. DMOG:SAR405 | -0.0002657 | No | ns | 0.2526 |
| Vehicle:SAR405 vs. DMOG:Vehicle | -0.001209 | Yes | **** | <0.0001 |
| Vehicle:SAR405 vs. DMOG:SAR405 | -0.0003138 | No | ns | 0.1978 |
| DMOG:Vehicle vs. DMOG:SAR405 | 0.0008950 | Yes | **** | <0.0001 |

**Figure S4.** DMOG induced mitophagy requires macroautophagy in both ubi:mitoKeima and ubi:mitoGR larvae. (A) Full statistical analyses of the experiment shown in Fig. 8A. Ubi:mitoKeima larvae treated with 100 $\mu$M DMOG and 20 $\mu$M SAR405 or the associated vehicles from 3-4 dpf and analyzed for Mitophagy Index. Data shown are median, 1$^{st}$ and 3$^{rd}$ quartiles, minimum and maximum. Two-way ANOVA with Holm-Sidak's multiple comparisons test. (A') Replication and full statistical analysis of experiment shown in Fig. 8A. Data shown are mean + SEM. *n* indicates number of larvae. Two-way ANOVA with Holm-Sidak's multiple comparisons test. (*B*) Representative images of ubi:mito-GR larvae treated with 100 $\mu$M DMOG and 20 $\mu$M SAR405 or the associated vehicles from 3-4 dpf and analyzed for Mitophagy Index. Scale, 20 $\mu$m. Data shown are mean + SEM. *n* indicates number of larvae. Two-way ANOVA with Holm-Sidak's multiple comparisons test.

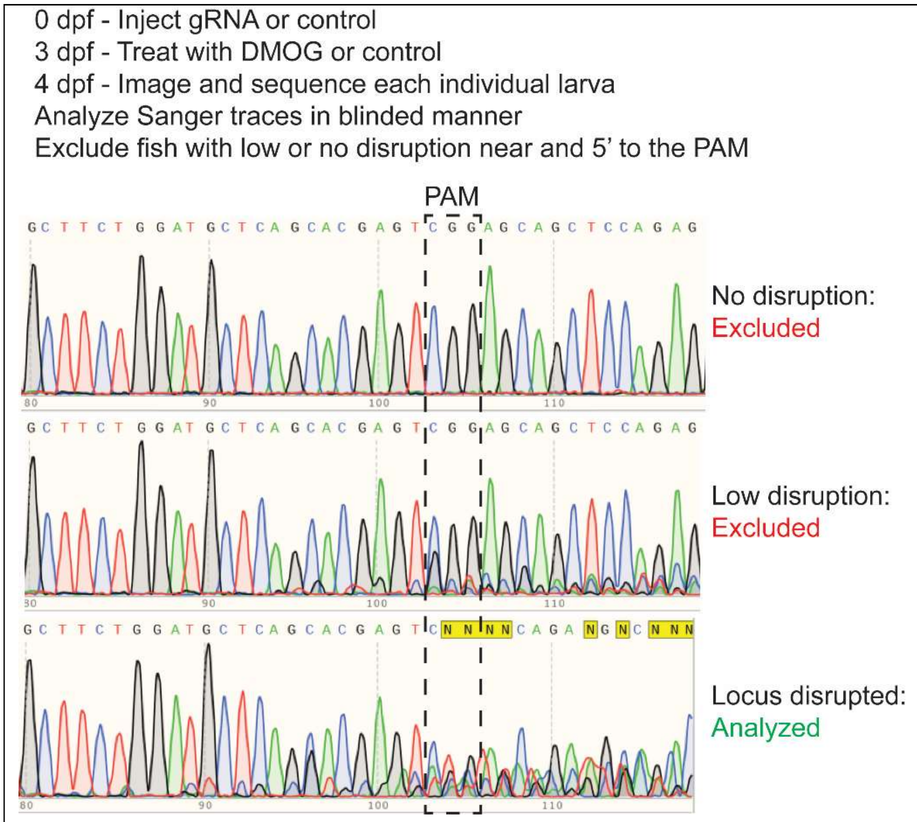


**Figure S5.** Experimental outline for high-efficiency CRISPR gene disruption experiments. 1-cell stage embryos were injected with CRISPR reagents or controls and raised until 70 hpf. Then the larvae were treated with DMOG or vehicle control until 98 hpf and imaged. Sequence analysis was performed on individual animals in a blinded fashion, and traces were binned into categories. “No disruption” indicates embryos with completely failed CRISPR possibly due to poor injections, and “Low disruption” indicates embryos with some locus disruption but a low level of mosaicism with the majority of cells containing the WT allele—these were removed from analysis. “Locus disrupted” indicates embryos with traces where the WT sequence could not be distinguished from edited sequences, and these were included in analyses.

**A**

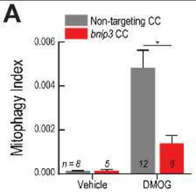


### Same as Fig. 7B

| Holm-Sidak's multiple comparisons test | Predicted (LS) mean diff. | Significant? | Summary | Adjusted P Value |
|---|---|---|---|---|
| Vehicle:Non-targeting vs. Vehicle:bnip3 CC | -1.575e-005 | No | ns | 0.9880 |
| Vehicle:Non-targeting vs. DMOG:Non-targeting | -0.004687 | Yes | **** | <0.0001 |
| Vehicle:Non-targeting vs. DMOG:bnip3 CC | -0.001259 | No | ns | 0.5062 |
| Vehicle:bnip3 cc1 vs. DMOG:Non-targeting | -0.004671 | Yes | *** | 0.0002 |
| Vehicle:bnip3 cc1 vs. DMOG:bnip3 CC | -0.001243 | No | ns | 0.5062 |
| **DMOG:Non-targeting vs. DMOG:bnip3 CC** | **0.003428** | **Yes** | ** | **0.0031** |

**A’**

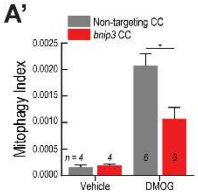


### Repeat of Fig. 7B

| Holm-Sidak's multiple comparisons test | Predicted (LS) mean diff. | Significant? | Summary | Adjusted P Value |
|---|---|---|---|---|
| Vehicle:Non-targeting vs. Vehicle:bnip3 CC | -3.051e-005 | No | ns | 0.9203 |
| Vehicle:Non-targeting vs. DMOG:Non-targeting | -0.001921 | Yes | **** | <0.0001 |
| Vehicle:Non-targeting vs. DMOG:bnip3 CC | -0.0009109 | Yes | * | 0.0128 |
| Vehicle:bnip3 CC vs. DMOG:Non-targeting | -0.001890 | Yes | **** | <0.0001 |
| Vehicle:bnip3 CC vs. DMOG:bnip3 CC | -0.0008804 | Yes | * | 0.0128 |
| **DMOG:Non-targeting vs. DMOG:bnip3 CC** | **0.001010** | **Yes** | ** | **0.0032** |

**B**

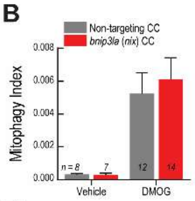


### Same as Fig. 7C

| Holm-Sidak's multiple comparisons test | Predicted (LS) mean diff. | Significant? | Summary | Adjusted P Value |
|---|---|---|---|---|
| Vehicle:Non-targeting vs. Vehicle:bnip3la (nix) CC | 2.339e-005 | No | ns | 0.9905 |
| Vehicle:Non-targeting vs. DMOG:Non-targeting | -0.004929 | Yes | * | 0.0262 |
| Vehicle:Non-targeting vs. DMOG:bnip3la (nix) CC | -0.005797 | Yes | ** | 0.0077 |
| Vehicle:bnip3la (nix) CC vs. DMOG:Non-targeting | -0.004952 | Yes | * | 0.0262 |
| Vehicle:bnip3la (nix) CC vs. DMOG:bnip3la (nix) CC | -0.005820 | Yes | ** | 0.0093 |
| DMOG:Non-targeting vs. DMOG:bnip3la (nix) CC | -0.0008679 | No | ns | 0.8065 |

**B’**

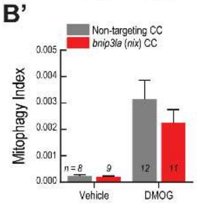


### Repeat of Fig. 7C

| Holm-Sidak's multiple comparisons test | Predicted (LS) mean diff. | Significant? | Summary | Adjusted P Value |
|---|---|---|---|---|
| Vehicle:Non-targeting vs. Vehicle:bnip3la (nix) CC | 2.903e-005 | No | ns | 0.9715 |
| Vehicle:Non-targeting vs. DMOG:Non-targeting | -0.002906 | Yes | ** | 0.0024 |
| Vehicle:Non-targeting vs. DMOG:bnip3la (nix) CC | -0.002007 | Yes | * | 0.0396 |
| Vehicle:bnip3la (nix) CC vs. DMOG:Non-targeting | -0.002935 | Yes | ** | 0.0018 |
| Vehicle:bnip3la (nix) CC vs. DMOG:bnip3la (nix) | -0.002036 | Yes | * | 0.0386 |
| DMOG:Non-targeting vs. DMOG:bnip3la (nix) CC | 0.0008993 | No | ns | 0.3643 |

**C**

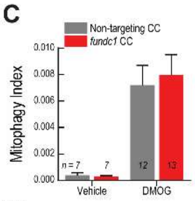


### Same as Fig. 7D

| Holm-Sidak's multiple comparisons test | Predicted (LS) mean diff. | Significant? | Summary | Adjusted P Value |
|---|---|---|---|---|
| Vehicle:Non-targeting vs. Vehicle:fundc1 CC | 9.286e-005 | No | ns | 0.9684 |
| Vehicle:Non-targeting vs. DMOG:Non-targeting | -0.006803 | Yes | ** | 0.0083 |
| Vehicle:Non-targeting vs. DMOG:fundc1 CC | -0.007582 | Yes | ** | 0.0038 |
| Vehicle:fundc1 CC vs. DMOG:Non-targeting | -0.006896 | Yes | ** | 0.0083 |
| Vehicle:fundc1 CC vs. DMOG:fundc1 CC | -0.007675 | Yes | ** | 0.0038 |
| DMOG:Non-targeting vs. DMOG:fundc1 CC | -0.0007790 | No | ns | 0.8831 |

**C’**

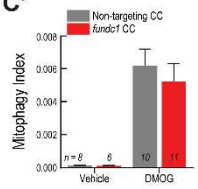


### Repeat of Fig. 7D

| Holm-Sidak's multiple comparisons test | Predicted (LS) mean diff. | Significant? | Summary | Adjusted P Value |
|---|---|---|---|---|
| Vehicle:Nontargeting vs. Vehicle:fundc1 CC | 1.417e-005 | No | ns | 0.9923 |
| Vehicle:Nontargeting vs. DMOG:Nontargeting | -0.006057 | Yes | *** | 0.0003 |
| Vehicle:Nontargeting vs. DMOG:fundc1 CC | -0.005108 | Yes | ** | 0.0011 |
| Vehicle:fundc1 CC vs. DMOG:Nontargeting | -0.006071 | Yes | *** | 0.0006 |
| Vehicle:fundc1 CC vs. DMOG:fundc1 CC | -0.005122 | Yes | ** | 0.0022 |
| DMOG:Nontargeting vs. DMOG:fundc1 CC | 0.0009490 | No | ns | 0.6697 |

**D**

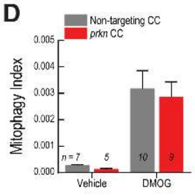


### Same as Fig. 7F

| Holm-Sidak's multiple comparisons test | Predicted (LS) mean diff. | Significant? | Summary | Adjusted P Value |
|---|---|---|---|---|
| Vehicle:Non-targeting vs. Vehicle:prkn CC | 0.0001417 | No | ns | 0.8886 |
| Vehicle:Non-targeting vs. DMOG:Non-targeting | -0.002903 | Yes | ** | 0.0038 |
| Vehicle:Non-targeting vs. DMOG:prkn CC | -0.002598 | Yes | ** | 0.0089 |
| Vehicle:prkn CC vs. DMOG:Non-targeting | -0.003045 | Yes | ** | 0.0056 |
| Vehicle:prkn CC vs. DMOG:prkn CC | -0.002739 | Yes | ** | 0.0100 |
| DMOG:Non-targeting vs. DMOG:prkn CC | 0.0003057 | No | ns | 0.8886 |

**D’**

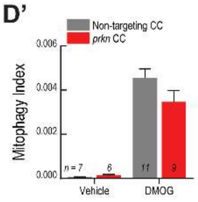


### Repeat of Fig. 7F

| Holm-Sidak's multiple comparisons test | Predicted (LS) mean diff. | Significant? | Summary | Adjusted P Value |
|---|---|---|---|---|
| Vehicle:Non-targeting vs. Vehicle:prkn CC | -0.0001774 | No | ns | 0.9106 |
| Vehicle:Non-targeting vs. DMOG:Non-targeting | -0.01121 | Yes | **** | <0.0001 |
| Vehicle:Non-targeting vs. DMOG:prkn CC | -0.008520 | Yes | **** | <0.0001 |
| Vehicle:prkn CC vs. DMOG:Non-targeting | -0.01104 | Yes | **** | <0.0001 |
| Vehicle:prkn CC vs. DMOG:prkn | -0.008343 | Yes | **** | <0.0001 |
| DMOG:Non-targeting vs. DMOG:prkn CC | 0.002693 | No | ns | 0.0820 |

**Figure S6.** Replication and full statistical analyses for Fig. 8. (*A*) Full statistical analysis for experiment shown in Fig. 8B. (*A’*) Replication and full statistical analysis of experiment shown in Fig. 8B. (*B*) Full statistical analysis for experiment shown in Fig. 8C. (*B'*) Replication and full statistical analysis of experiment shown in Fig. 8C. (*C*) Full statistical analysis for experiment shown in Fig. 8D. (*C'*) Replication and full statistical analysis of experiment shown in Fig. 8D. (*D*) Full statistical analysis for experiment shown in Fig. 8F. (*D'*) Replication and full statistical analysis of experiment shown in Fig. 8F. (*A*-*D*) Data shown are mean + SEM. *n* indicates number of larvae. Two-way ANOVA with Holm-Sidak's multiple comparisons test.

**A**

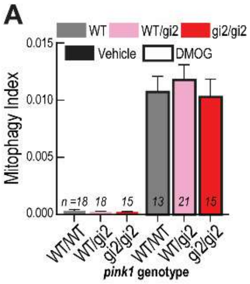


**Same as Fig. 7E**

| *Holm-Sidak's multiple comparisons test* | *Predicted (LS) mean diff.* | *Significant?* | *Summary* | *Adjusted P Value* |
|---|---|---|---|---|
| WT/WT:Vehicle vs. WT/WT:DMOG | -0.01039 | Yes | **** | <0.0001 |
| WT/WT:Vehicle vs. WT/gi2:Vehicle | 7.956e-005 | No | ns | 0.9998 |
| WT/WT:Vehicle vs. WT/gi2:DMOG | -0.01145 | Yes | **** | <0.0001 |
| WT/WT:Vehicle vs. gi2/gi2:Vehicle | 0.0001079 | No | ns | 0.9998 |
| WT/WT:Vehicle vs. gi2/gi2:DMOG | -0.009954 | Yes | **** | <0.0001 |
| WT/WT:DMOG vs. WT/gi2:Vehicle | 0.01047 | Yes | **** | <0.0001 |
| WT/WT:DMOG vs. WT/gi2:DMOG | -0.001061 | No | ns | 0.9543 |
| WT/WT:DMOG vs. gi2/gi2:Vehicle | 0.01050 | Yes | **** | <0.0001 |
| WT/WT:DMOG vs. gi2/gi2:DMOG | 0.0004354 | No | ns | 0.9976 |
| WT/gi2:Vehicle vs. WT/gi2:DMOG | -0.01153 | Yes | **** | <0.0001 |
| WT/gi2:Vehicle vs. gi2/gi2:Vehicle | 2.836e-005 | No | ns | 0.9998 |
| WT/gi2:Vehicle vs. gi2/gi2:DMOG | -0.01003 | Yes | **** | <0.0001 |
| WT/gi2:DMOG vs. gi2/gi2:Vehicle | 0.01156 | Yes | **** | <0.0001 |
| WT/gi2:DMOG vs. gi2/gi2:DMOG | 0.001496 | No | ns | 0.8585 |
| gi2/gi2:Vehicle vs. gi2/gi2:DMOG | -0.01006 | Yes | **** | <0.0001 |

**B**

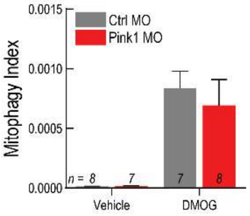


| *Holm-Sidak's multiple comparisons test* | *Predicted (LS) mean diff.* | *Significant?* | *Summary* | *Adjusted P Value* |
|---|---|---|---|---|
| Vehicle:Ctrl MO vs. Vehicle:Pink1 MO | -7.666e-007 | No | ns | 0.9968 |
| Vehicle:Ctrl MO vs. DMOG:Ctrl MO | -0.0008222 | Yes | ** | 0.0012 |
| Vehicle:Ctrl MO vs. DMOG:Pink1 MO | -0.0006803 | Yes | ** | 0.0041 |
| Vehicle:Pink1 MO vs. DMOG:Ctrl MO | -0.0008215 | Yes | ** | 0.0015 |
| Vehicle:Pink1 MO vs. DMOG:Pink1 MO | -0.0006795 | Yes | ** | 0.0043 |
| DMOG:Ctrl MO vs. DMOG:Pink1 MO | 0.0001420 | No | ns | 0.7113 |

**Figure S7.** *pink1* depletion has no effect on DMOG-induced mitophagy. (A) This figure is the same data as Figure 8E showing full statistical analyses. (B) Mitophagy Index quantification of 72 hpf larvae injected with a control (Ctrl) MO or *pink1* MO and treated with vehicle control (DMSO) or DMOG from 48-72 hpf. Data shown are mean + SEM. *n* indicates number of larvae. ns, not significant (*P*>0.05, two-way ANOVA with Holm-Sidak's correction). Comparisons between all Vehicle and DMOG-treated samples were significant (*P*<0.05.).

A

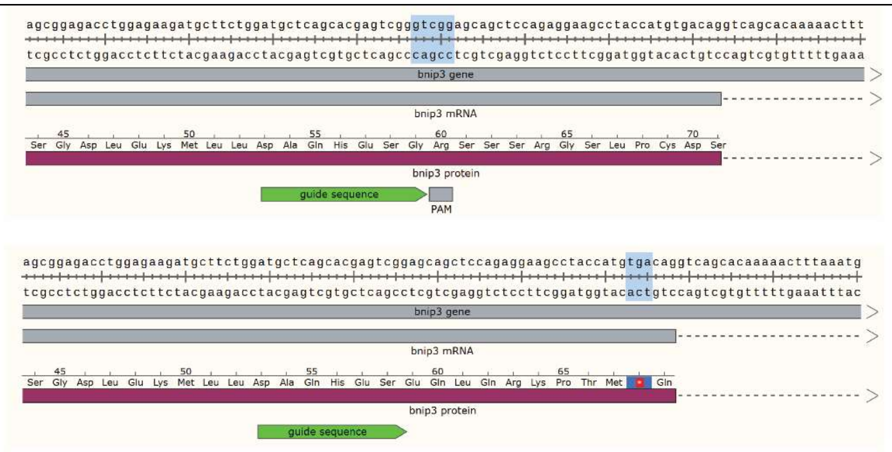


C

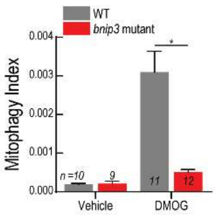


**Same as Fig. 7G**

| *Holm-Sidak's multiple comparisons test* | *Predicted (LS) mean diff.* | *Significant?* | *Summary* | *Adjusted P Value* |
|---|---|---|---|---|
| Vehicle:WT vs. Vehicle:bnip3-/- | -2.522e-005 | No | ns | 0.9533 |
| Vehicle:WT vs. DMOG:WT | -0.002908 | Yes | **** | <0.0001 |
| Vehicle:WT vs. DMOG:bnip3-/- | -0.0003213 | No | ns | 0.8100 |
| Vehicle:bnip3-/- vs. DMOG:WT | -0.002883 | Yes | **** | <0.0001 |
| Vehicle:bnip3-/- vs. DMOG:bnip3-/- | -0.0002961 | No | ns | 0.8100 |
| **DMOG:WT vs. DMOG:bnip3-/-** | **0.002587** | **Yes** | ******** | **<0.0001** |

C’

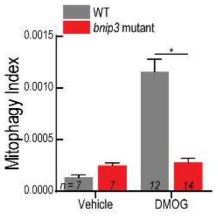


**Repeat of Fig. 7G**

| *Holm-Sidak's multiple comparisons test* | *Predicted (LS) mean diff.* | *Significant?* | *Summary* | *Adjusted P Value* |
|---|---|---|---|---|
| Vehicle:WT vs. Vehicle:bnip3-/- | -0.0001165 | No | ns | 0.6353 |
| Vehicle:WT vs. DMOG:WT | -0.001022 | Yes | **** | <0.0001 |
| Vehicle:WT vs. DMOG:bnip3-/- | -0.0001481 | No | ns | 0.5175 |
| Vehicle:bnip3-/- vs. DMOG:WT | -0.0009055 | Yes | **** | <0.0001 |
| Vehicle:bnip3-/- vs. DMOG:bnip3-/- | -3.155e-005 | No | ns | 0.7898 |
| **DMOG:WT vs. DMOG:bnip3-/-** | **0.0008739** | **Yes** | ******** | **<0.0001** |

**Figure S8.** Mutation of *bnip3* mitigates Hif-induced mitophagy. (A) 3’ portion of exon 2 in the WT *bnip3* allele with CRISPR guide sequencing and protospacer adjacent motif (PAM) labeled. Blue highlight shows the 5-bp that were deleted in the mutant allele. (B) Sequence of the 3’ portion of exon 2 in the *bnip3* mutant allele. Blue highlight shows the premature stop codon. (C) Full statistical analysis for experiment show in Fig. 8G. (C’) Replication and full statistical analysis of experiment show in the Fig. 8G. (C,C’) Data shown are mean + SEM. *n* indicates number of larvae. Two-way ANOVA with Holm-Sidak’s multiple comparisons test.

### Movie Legends

**Movie S1.** Supports Figures 3A and 3B: depicts the large 3D data sets acquired and the segmentation and analysis method employed to calculate Mitophagy Index using a representative 7 dpf fasted ubi:mitoKeima larva. Keima440 is in green, Keima561 is in magenta. At the beginning of the movie, a 3D rendered maximum intensity projection is displayed and rotated. Then, the ratiometric image derived from Keima561/Keima440 is shown. Lastly, the isosurface rendering objects are shown. The magenta surface objects represent mitolysosomes and the green surface object represents the total tissue volume analyzed.

**Movie S2.** Supports Figure 4: *in vivo* mitophagy dynamics: mitophagy kinetics and mitolysosome fission. Healthy mitochondria (Keima440) appear green. Mitolysosomes (Keima561/440) appear magenta. Maximum intensity projections.

**Movie S3.** Supports Figure 5: mitophagy occurs piecemeal. Healthy mitochondria (Keima440) appear cyan. Mitolysosomes (Keima561/Keima440) are magenta. EGFP-Lc3 is yellow. Isosurface renderings.

### Supplemental References

Berger J & Currie PD (2013) 503Unc, a Small and Muscle-Specific Zebrafish Promoter. *Genesis* 51: 443–447

Choi TY, Ninov N, Stainier DYR & Shin D (2014) Extensive conversion of hepatic biliary epithelial cells to hepatocytes after near total loss of hepatocytes in zebrafish. *Gastroenterology* 146: 776–788

Don EK, Formella I, Badrock AP, Hall TE, Morsch M, Hortle E, Hogan A, Chow S, Gwee SSL, Stoddart JJ, *et al* (2017) A Tol2 Gateway-Compatible Toolbox for the Study of the Nervous System and Neurodegenerative Disease. *Zebrafish* 14: 69–72

Elks PM, Van Eeden FJ, Dixon G, Wang X, Reyes-Aldasoro CC, Ingham PW, Whyte MKB, Walmsley SR & Renshaw SA (2011) Activation of hypoxia-inducible factor-1α (hif-1α) delays inflammation resolution by reducing neutrophil apoptosis and reverse migration in a zebrafish inflammation model. *Blood* 118: 712–722

Harris JM, Esain V, Frechette GM, Harris LJ, Cox AG, Cortes M, Garnaas MK, Carroll KJ,

Cutting CC, Khan T, *et al* (2013) Glucose metabolism impacts the spatiotemporal onset and magnitude of HSC induction in vivo. *Blood* 121: 2483–2493

He C, Bartholomew CR, Zhou W & Klionsky DJ (2009) Assaying autophagic activity in transgenic GFP-Lc3 and GFP-Gabarap zebrafish embryos. *Autophagy* 5: 520–526

Katayama H, Kogure T, Mizushima N, Yoshimori T & Miyawaki A (2011) A sensitive and quantitative technique for detecting autophagic events based on lysosomal delivery. *Chem Biol* 18: 1042–1052

Kwan KM, Fujimoto E, Grabher C, Mangum BD, Hardy ME, Campbell DS, Parant JM, Yost HJ, Kanki JP & Chien C Bin (2007) The Tol2kit: A multisite gateway-based construction Kit for Tol2 transposon transgenesis constructs. *Dev Dyn* 236: 3088–3099

Lee E, Koo Y, Ng A, Wei Y, Luby-Phelps K, Juraszek A, Xavier RJ, Cleaver O, Levine B & Amatruda JF (2014) Autophagy is essential for cardiac morphogenesis during vertebrate development. *Autophagy* 10: 572–587

Lee E, Wei Y, Zou Z, Tucker K, Rakheja D, Levine B & Amatruda JF (2016) Genetic inhibition of autophagy promotes p53 loss-of-heterozygosity and tumorigenesis. *Oncotarget* 7

Mosimann C, Kaufman CK, Li P, Pugach EK, Tamplin OJ & Zon LI (2011) Ubiquitous transgene expression and Cre-based recombination driven by the ubiquitin promoter in zebrafish. *Development* 138: 169–177

Priyadarshini M, Tuimala J, Chen YC & Panula P (2013) A zebrafish model of PINK1 deficiency reveals key pathway dysfunction including HIF signaling. *Neurobiol Dis* 54: 127–138

Rojansky R, Cha MY & Chan DC (2016) Elimination of paternal mitochondria in mouse embryos occurs through autophagic degradation dependent on PARKIN and MUL1. *Elife* 5: e17896
